# MCell4 with BioNetGen: A Monte Carlo Simulator of Rule-Based Reaction-Diffusion Systems with Python Interface

**DOI:** 10.1101/2022.05.17.492333

**Authors:** Adam Husar, Mariam Ordyan, Guadalupe C. Garcia, Joel G. Yancey, Ali S. Saglam, James R. Faeder, Thomas M. Bartol, Mary B. Kennedy, Terrence J. Sejnowski

## Abstract

Biochemical signaling pathways in living cells are often highly organized into spatially segregated volumes, membranes, scaffolds, subcellular compartments, and organelles comprising small numbers of interacting molecules. At this level of granularity stochastic behavior dominates, well-mixed continuum approximations based on concentrations break down and a particle-based approach is more accurate and more efficient. We describe and validate a new version of the open-source MCell simulation program (MCell4), which supports generalized 3D Monte Carlo modeling of diffusion and chemical reaction of discrete molecules and macromolecular complexes in solution, on surfaces representing membranes, and combinations thereof. The main improvements in MCell4 compared to the previous versions, MCell3 and MCell3-R, include a Python interface and native BioNetGen reaction language (BNGL) support. MCell4’s Python interface opens up completely new possibilities for interfacing with external simulators to allow creation of sophisticated event-driven multiscale/multiphysics simulations. The native BNGL support, implemented through a new open-source library libBNG (also introduced in this paper), provides the capability to run a given BNGL model spatially resolved in MCell4 and, with appropriate simplifying assumptions, also in the BioNetGen simulation environment, greatly accelerating and simplifying model validation and comparison.

## 1 Introduction

Living cells are complex structures in which biomolecules and biochemical processes are spatially organized and span the extracellular space, plasma membrane, cytosol and subcellular organelles. These biochemical processes are intrinsically multiscale in nature because they are based on molecular interactions on a small scale leading to emergent behavior of cells on a larger scale. Becajuse of the dynamic nature of biochemical processes on different temporal and spatial scales, appropriate mathematical tools are required to understand the underlying dynamics and to dissect the mechanisms that control system behavior [1]. Overall, understanding how cellular design dictates function is essential to understanding health and disease in the brain, heart, and elsewhere. MCell (Monte Carlo Cell) is a biochemistry simulation tool that uses spatially realistic 3D cellular models and stochastic Monte Carlo algorithms to simulate the movements and interactions of discrete molecules within and between cells[2, 3, 4, 5]. Here we describe MCell4, a new version of MCell.

One of the most important new features in MCell4 is a flexible Python application programming interface (API) that allows coupling between MCell and other simulation engines or other custom code. By itself MCell performs particle-based reaction-diffusion simulations on spatial and temporal scales from nm to µm and from µs to 10s of seconds. MCell4’s Python API extends its capabilities by facilitating the generation of multiscale hybrid models, as we demonstrate here with an example.

A second important addition to MCell4 is efficient support for rule-based modeling by making use of the BioNetGen (BNG) Language (BNGL). BNG is an open source software package for representing and simulating biochemical reactions [6]. Although powerful, BNG models are non-spatial. Support for models implemented in BNGL within MCell4 permits determination of the role of space in different reaction scenarios. This is not a trivial task because the time scales of diffusion and of reactions [7], as well as the spatial localization of proteins, influence the results.

We first present the design principles of MCell4 and its API, and next we introduce the new BioNetGen library. Finally we demonstrate some of the new features in MCell 4 with examples and present a hybrid model that couples spatial simulations in MCell with ordinary differential equations (ODEs).

### 1.1 Particle-Based Reaction Dynamics Tools

In particle-based reaction-diffusion simulations, each molecule is represented as an individual agent. Molecules diffuse either within volumes or on membrane surfaces and may affect each other by reacting upon collision. A review of currently maintained particle-based stochastic simulators which describes Smoldyn [7], eGFRD [8], SpringSaLaD [9], ReaDDy [10], and MCell3 was recently published in [11].

MCell is a particle-based simulator that represents volume molecules as point particles and surface molecules as area-filling tiles on surfaces. The typical simulation time-step in MCell is 1 µs, and the simulated times can stretch from milliseconds to minutes. Briefly, MCell operates as follows. As a volume molecule diffuses through space by random Brownian motion, all volume molecules within a given radius (i.e. the interaction radius, *r_int_*) along its trajectory, or the single surface molecule located at the point of collision on a surface, are considered as possible reaction partners. As a surface molecule diffuses it is first moved to its final position after one time step and any surface molecules immediately adjacent to that final position are considered as possible reaction partners. Molecules diffusing in 3D volumes do not themselves have volume (i.e. no volume exclusion). The collision cross-section area for interactions among volume molecules is derived from *r_int_*. Molecules on membrane surfaces occupy a fixed area defined by the individual triangular grid elements (tiles) created by subdividing the surface mesh triangles with a barycentric grid. The collision cross-section for interactions between volume and surface molecules and among surface molecules is derived from the density of the barycentric surface grid. MCell is able to represent arbitrary geometries comprised of triangulated surface meshes. Thus complex models such as a 180 µm^3^ 3 dimensional serial electron microscopic reconstruction of hippocampal neuropil have been used to construct a geometrically-precise and biophysically accurate simulation of synaptic transmission and calcium dynamics in neuronal synapses [5]. A detailed description of the mathematical foundations of MCell’s algorithms can be found in these references [2, 3, 4].

MCell3-R [12], a precursor of of MCell4, is an extension of MCell that supports BNGL [13] and allows modeling of protein complexes or polymers by using rule-based definition of reactions. MCell3-R uses a library called NFSim [14] to compute the products of reaction rules for reactions described in BNGL.

MCell4 is an entirely new implementation of MCell, written in C++. It provides a versatile Python interface in addition to many other improvements. In particular it runs significantly faster when simulating complex reaction networks expressed as rules in BNGL. And most of the features of MCell that were introduced previously [4] have been retained. Here we briefly describe the motivations for introducing new features in MCell4.

### 1.2 Motivation for the MCell4 Python Application Programming Interface

We had two important motivations for the creation of the MCell4 Python API: 1) to give the users the freedom to customize their models in a full-featured modern programming language, and 2) to create an easy way to couple MCell4 with other simulation platforms to allow multi-scale, multi-physics simulations.

The main goal when designing the new API for MCell4 was to allow definition of complex models combining many reaction pathways distributed over complex geometry. Thus, a main requirement was to enable modularity with reusable components that can be independently validated. With this feature one can build complex models by combining existing modules with new ones.

As in the approach in the PySB modeling framework [15], a model in MCell4 is seen as a piece of software, allowing the same processes used in software development to be applied to biological model development. The most important such processes are: 1) incremental development where the model is built step by step, relying on solid foundations of modeling that has been validated previously, 2) modularity that provides the capability to create self-contained, reusable libraries, 3) unit testing and validation to verify that parts of the model behave as expected, and 4) human-readable and writable model code that can be stored with git or other code version control software. In addition to being essential for incremental development, this also allows code reviews [16] so that other team members can inspect the latest changes to the model and can contribute their own modules to the growing code base.

### 1.3 Motivation for a New BioNetGen Library

NFSim [14] is a C++ library that provides BioNetGen support, implements the network-free method, and is used in MCell3-R [12]. To use a BNGL model in MCell3-R, the BNGL file first needs to be parsed by the BioNetGen compiler; then, a converter generates a file containing MCell Model Description Language (MDL), a file with rule-based extensions to MDL (MDLR), and additional XML files required by the NFSim library. These files then constitute the model for simulation in MCell3-R. The disadvantage of this approach is that the original BNGL file is no longer present in the MCell3-R model. Thus each time changes are made to the model, the converter must be run again, and any changes made by hand to the MCell3-R model files will be lost. MCell3-R also has performance and memory consumption problems when the simulated system has a large number of potential reactions.

To create a seamless integration of BNGL with MCell4 we implemented a new library for the BioNetGen language that contains a BNGL parser and a network-free BNG reaction engine the main purpose of which is to compute reaction products for a given set of reaction rules and reactants. This BNG library (libBNG) was designed to be independent of MCell4 in mind so that it can be used in other simulation tools. libBNG does not yet support all of the special features and keywords of the BioNetGen tool suite. Most notably, BNGL functions are not supported, however the set of supported features is sufficient for any MCell4 model. And when a special function is needed, it can be represented in Python code with the MCell4 API. The source code of libBNG is available under the MIT license in Reference [17].

### 1.4 Features of MCell4

Here we briefly describe some of the features of MCell4. In the results section we present a few relevant examples specifically to demonstrate some of these features. We indicate which example illustrates the mentioned feature.

#### 1.4.1 Python/C++ API for Model Creation and Execution

All models can now be created in Python. CellBlender (see section 1.5) is useful for creation of many relatively simple models without the need to write Python code by hand. However, more complicated customized models will need to include a custom Python script. While CellBlender provides for inclusion of custom python scripts, for simplicity and explanatory power, all the examples presented in the results section of this paper are written solely in Python.

#### 1.4.2 Reactions are Now Written in BNGL

In MCell4 the reaction language is BNGL [13]. Thus, MCell4 fully supports rule-based reactions and all models use this feature.

Most importantly, the support for BNGL and NFSim means that MCell4 performs direct, agent-based evaluation of reaction rules and thus enables spatially-resolved network-free simulations of interactions between and among volume and surface molecules. The CaMKII holoenzyme model in the results section 3.1.3, for example, would not be possible without the spatial network-free algorithms implemented in MCell4.

#### 1.4.3 Ability to Go Back and Forth between MCell4 and BNG Simulator Environments

The new BNG library [17] allows direct loading and parsing of a BNG model that can then be placed within a realistic 3D cellular geometry. This allows comparison of the results of non-spatial (simulated with BioNetGen solvers) and spatial (simulated with MCell4) implementations of the same BNG model. Two of the examples in the results section demonstrate this feature: SNARE (3.1.1), and CaMKII (3.1.3).

If the spatial features are found to be unimportant for a given model, and simulation speed is of more concern, the BNGL file can be run as a separate module with the BNG simulator. See section 2.4.3 for an example.

#### 1.4.4 Other Advanced Features

Among the more advanced features introduced in MCell4 is the ability to implement transcellular and transmembrane interactions that occur between surface molecules located on separate membranes. MCell4 also supports both coarse-grained and fine-grained customization of models by customizing the time-step customization and by introducing event-driven callbacks. Callbacks implement custom Python code that runs when a particular reaction occurs or when a collision occurs between a molecule and a wall. An example of the use of callbacks to implement release of neurotransmitter when a SNARE complex is activated is shown in section-3.1.2.

Finally, the new Python API supports the ability to create multi-scale multi-physics hybrid simulations that take advantage of all the existing Python packages. For an example of a hybrid model see section 3.3.

### 1.5 Model Creation and Visualization in CellBlender

CellBlender is a Blender [18] addon that supports creation, execution, analysis, and visualization of MCell4 models. CellBlender has been updated from its previous MCell3 version and includes several new features: automatic generation of well structured Python code from the CellBlender representations of complete MCell4 models; execution, analysis, and visualization of these models; and visualization of simulation data generated by simulations of externally created Python-only models. CellBlender offers an easy way to begin using MCell through built-in examples (Fig. 1 shows an example of a model of the Rat Neuromuscular Junction), and tutorials [19].

**Figure 1:**
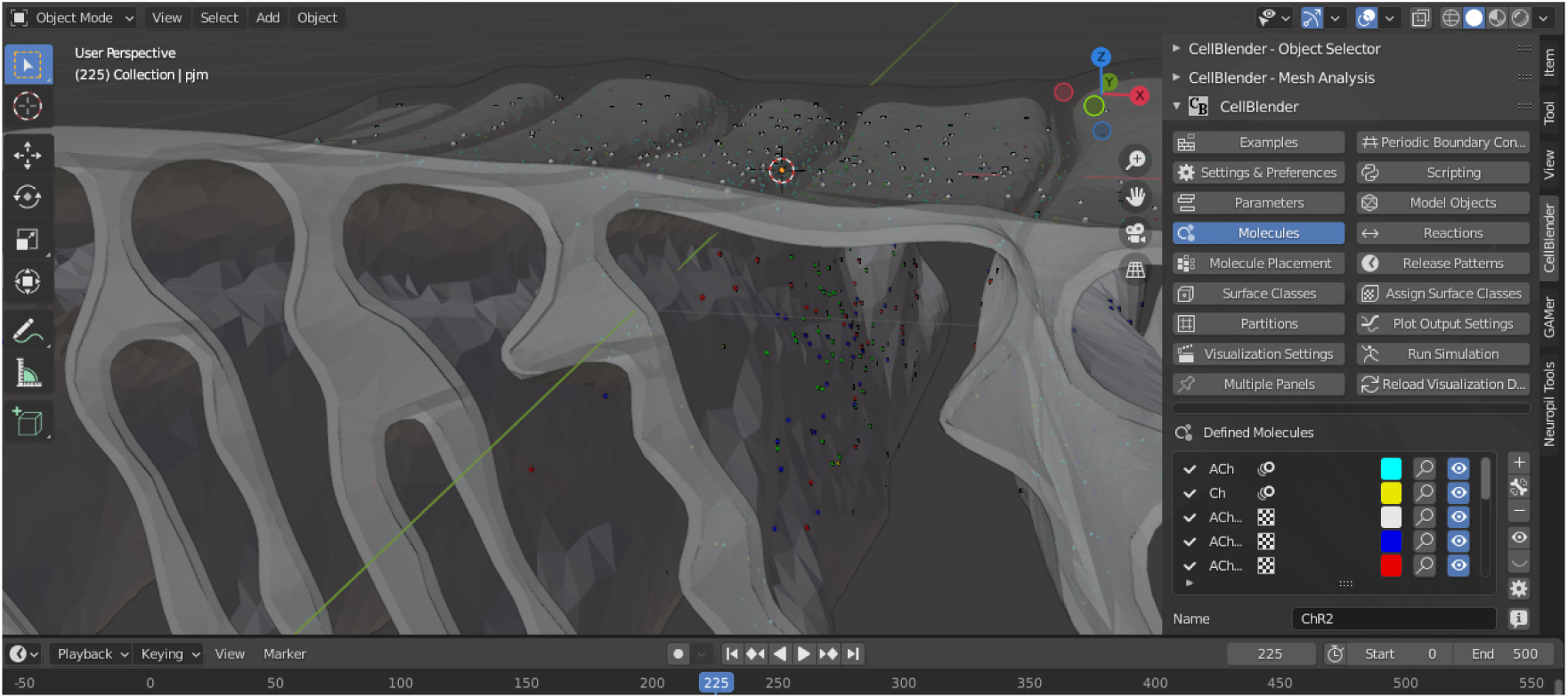
MCell4 models can be created, executed, and visualized using CellBlender, an addon for Blender. The capabilities of Blender are indispensable for creating complex geometries for MCell4 models.

## 2 Design and Implementation

### 2.1 MCell4: a Bird’s Eye View

We will briefly review MCell4’s architecture and fundamental aspects of its API, starting with Fig. 2.

**Figure 2:**
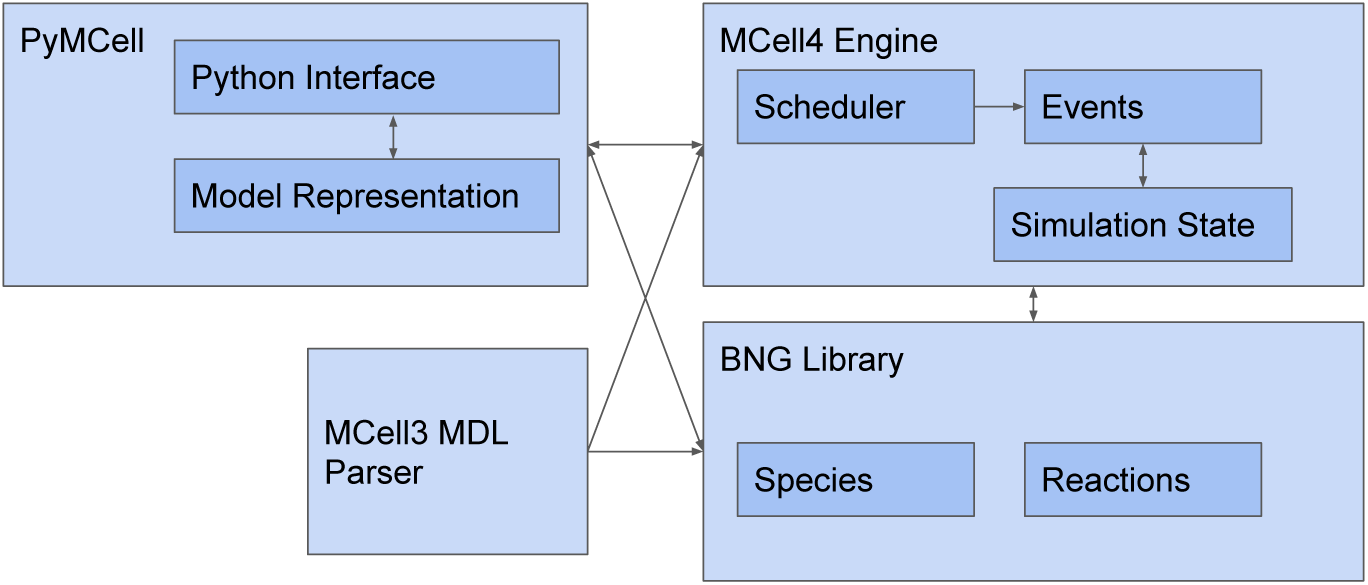
MCell4 is comprised of four main components: 1) The PyMCell library provides a Python interface and contains classes to hold the model representation, 2) The MCell4 engine implements the simulation algorithms, 3) The BNG (BioNetGen) library provides methods to resolve BioNetGen reactions, and 4) The MDL (Model Description Language) parser enables backwards compatibility with MCell3.

MCell simulations progress through time by a series of iterations. The duration of an iteration is given by a user-defined time step (usually 1 µs). The Scheduler keeps track of events to be run in each iteration. The main simulation loop implemented in the all-inclusive object called “World” requests the Scheduler to handle the all the events that occur in each iteration (Fig. 3) until the desired number of iterations have been completed.

**Figure 3:**
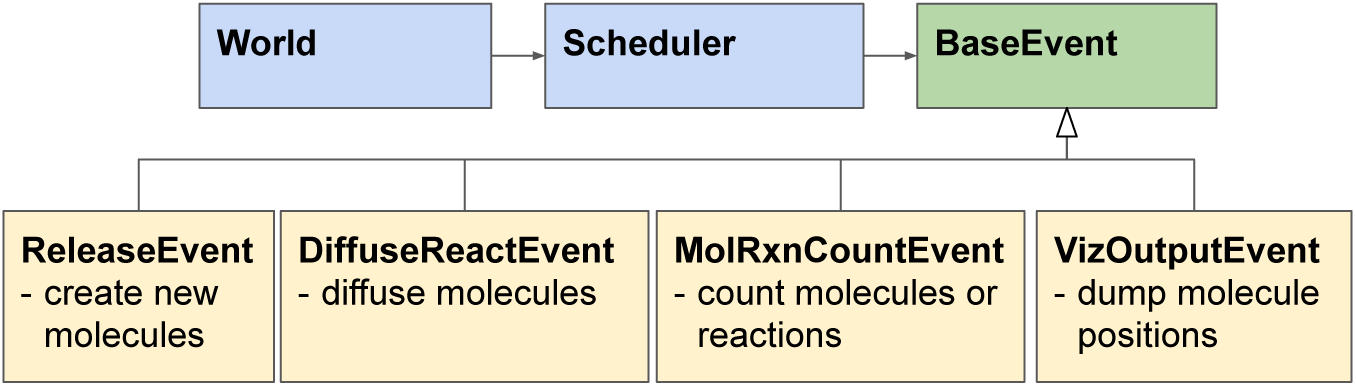
The Scheduler executes time step iterations which consist of discrete events executed in this order: 1) A ReleaseEvent creates new molecules, 2) A MolRxnCountEvent counts numbers of molecules or how many times a reaction occurrs, 3) A VizOutputEvent stores molecule locations for visualization in CellBlender, and 4) A DiffuseReactEvent implements diffusion of molecules, checks collisions, and executes reactions. Only the DiffuseReactEvent must be executed at each time step to move the time forward. The other events listed here are optional.

### 2.2 Python API Generator: A Closer Look

The MCell4 physics engine is implemented in C++. To ensure reliable correspondence between the representation of a model in Python and in C++, we have implemented a Python API generator which is used when building the MCell4 executable and Python module from source code. The API generator reads a high-level definition file in the YAML format and automatically generates all the base C++ classes, their *corresponding* Python API representations, code for informative error messages, and documentation. A consistent Python and C++ API contributes to the quality of the user experience when creating a model, and facilitates well maintained documentation.

The presence of the API generator, schematically represented in Fig. 4, ensures that when new features are added to MCell4, one only needs to modify a single API definition in the YAML format to ensure that both the API and the documentation reflect the new features.

**Figure 4:**
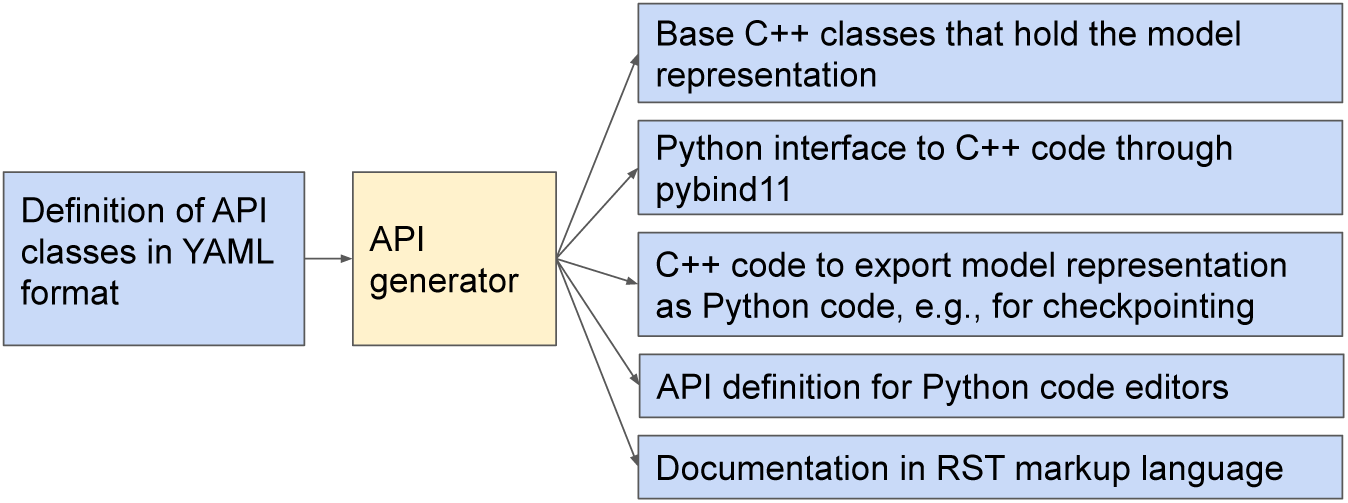
When MCell4 is built from its source code, the API generator reads a high-level definition of the MCell4 Python interface and generates code and documentation. Automatic generation of an API makes it possible to easily modify or extend the API while ensuring that all parts including documentation stay consistent. The API generator is a general tool that can also be used (with minor modifications) for other software tools that combine C++ and Python [20].

### 2.3 MCell4 Model Structure

A predefined model structure is important to enable reusability of model components (e.g., [21]). With a predefined model structure every piece of code for a given component (such as reaction definitions, geometry, initial model state, and observables) is in a file with a specified name and follows a predefined coding style. Such standardized model structure (shown in Fig. 5) aids in the reuse of code and simplifies creation of new models by leveraging existing model components. Another advantage of a predefined model structure is the capability to combine parts of existing models into one model (Fig. 6).

**Figure 5:**
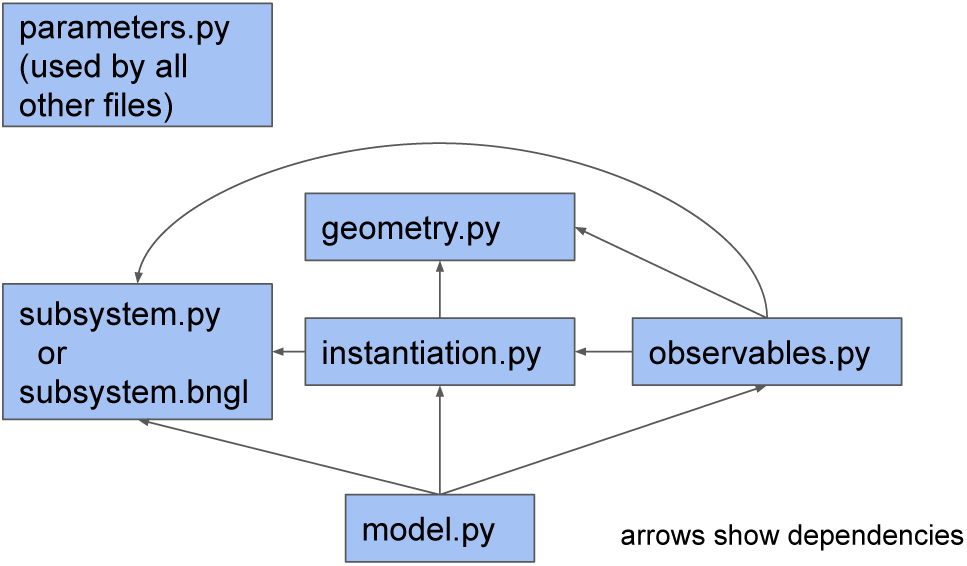
The main files included in a standard MCell4 model are: 1) parameters.py with all the model parameters, 2) subsystem.py that captures information on species and reactions in a way that is independent of a particular model and can be used as a reusable module, 3) geometry.py with a definition of 3D geometry objects, 4) instantiation.py that usually defines the initial model state, i.e., which geometry objects are created in the simulation and the number and positions of molecules to be released at a given time, 5) observables.py with lists of characteristics to be measured and saved in files during simulation, and 6) model.py in which all the parts of the model are assembled together and in which the the simulation loop with optional interactions with external simulators is defined. Model.py is the only required file.

**Figure 6:**
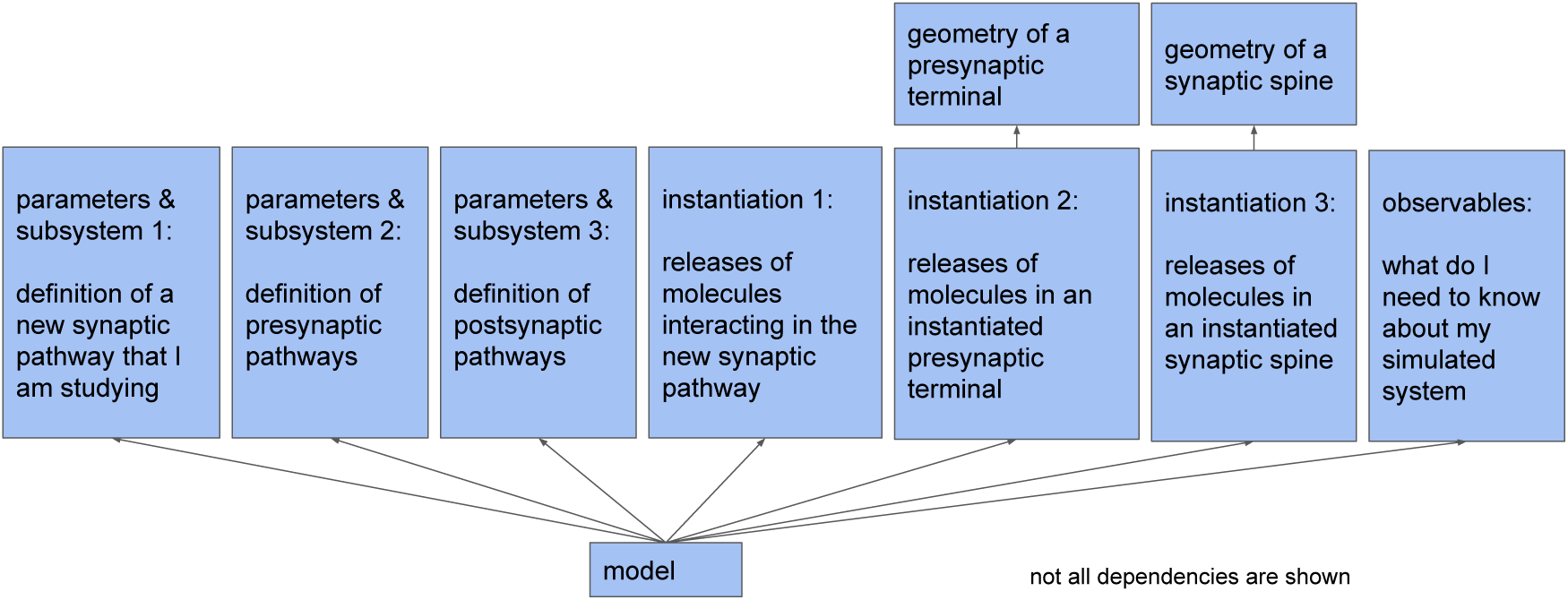
Modularity of a model allows assembly of multiple subsystem definitions into a single model. In the example shown here, individual modules are assembled to construct a model of a new synaptic pathway that is affected by other processes. The complete model includes modules that individually define the presynaptic terminal with its presynaptic pathways and the postsynaptic spine with its postsynaptic pathways.

#### 2.3.1 Example Model Using the MCell4 Python API

A simple example that shows the MCell4 API including Subsystem, Instantiation, and Model classes is shown in Fig. 7. Because of the simplicity of this example, we do not show the division into the separate files illustrated in Fig. 5.

**Figure 7:**
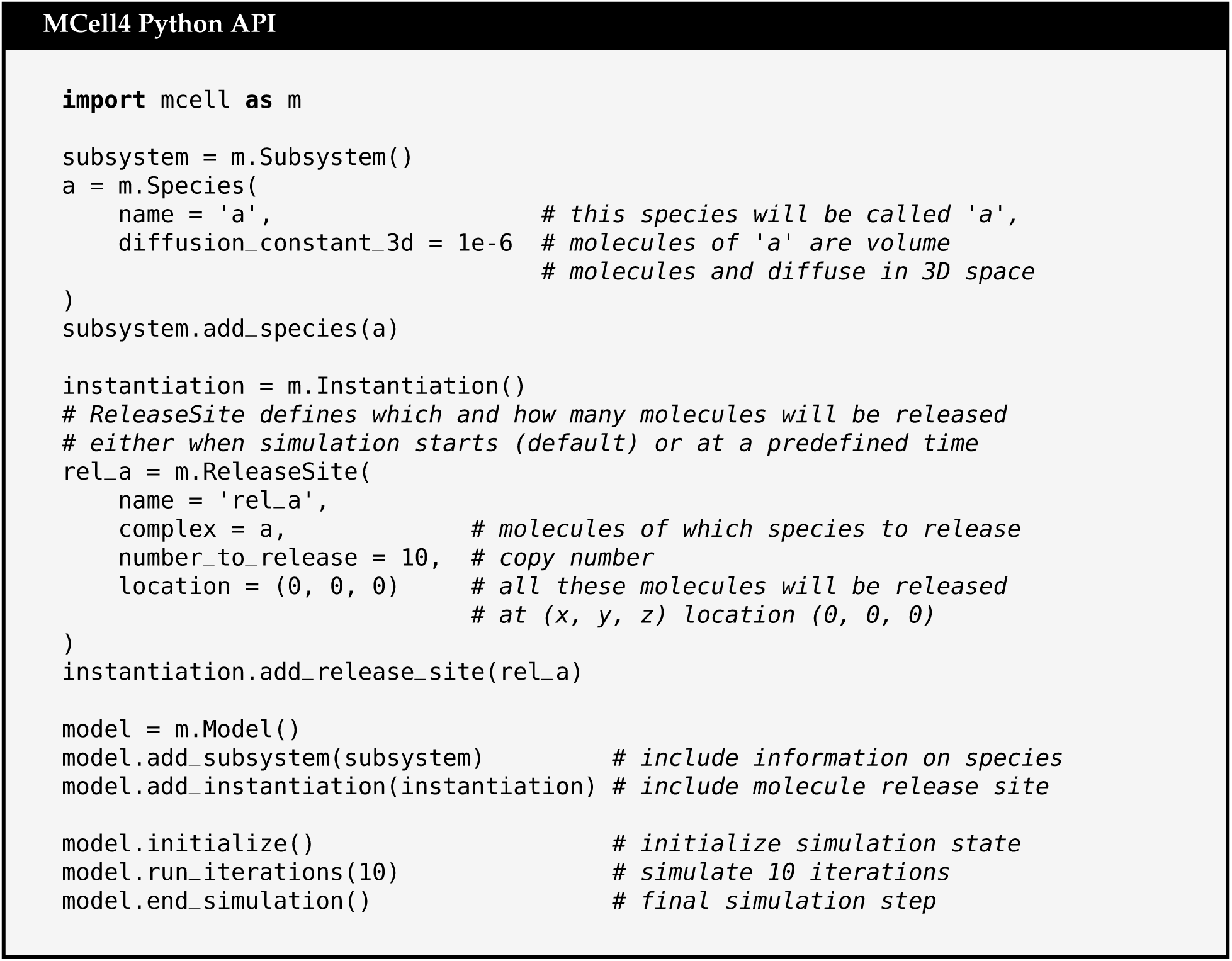
Example of a simple MCell4 model that releases 10 volume molecules of species ‘a’ and simulates their diffusion for 10 iterations with a default time step of 1 µs. Note that for this and following examples, a system variable, PYTHONPATH, must be set so that the Python interpreter knows where to find the MCell4 module [22]. Alternatively one can append to python’s search path from within the model file with the statement: import sys; sys.path.append(“/path/to/mcell4/libs”)

### 2.4 Graph-Based Approach To Protein Modeling

BNGL [23] supports intuitive modeling of protein complexes by representing them as undirected graphs. Such graphs contain two types of nodes: *elementary molecules* and *components*. Component nodes represent *binding sites* of the protein and can also express the *state* of the whole protein or of a binding site. A graph representing a *single protein* is implemented as an elementary molecule node with component nodes connected to it through *edges*. To form a dimer, two individual components of different proteins are bound by creating an edge between them. A graph with one or more elementary molecules with their components is called a *complex*. A *reaction rule* defines a graph transformation that manipulates the graph of reactants. A reaction rule usually manipulates edges to connect or disconnect complexes or change the state of a component. It can also modify the elementary molecules such as in the reaction A + B -> C where we do not care about the molecular details and do not need to model individual binding sites. An example of applying a reaction rule that connects complexes and changes the state is shown in Fig. 8. Note that what we call an “elementary molecule type” here is called a “molecule type” in BioNetGen. In MCell, “molecules” are defined as whole molecules such as protein complexes that act as individual agents in the simulation. For better clarity, we adopt the name “elementary molecule” for the base building blocks of complexes. The tool SpringSaLaD [9] uses the same distinction.

**Figure 8:**
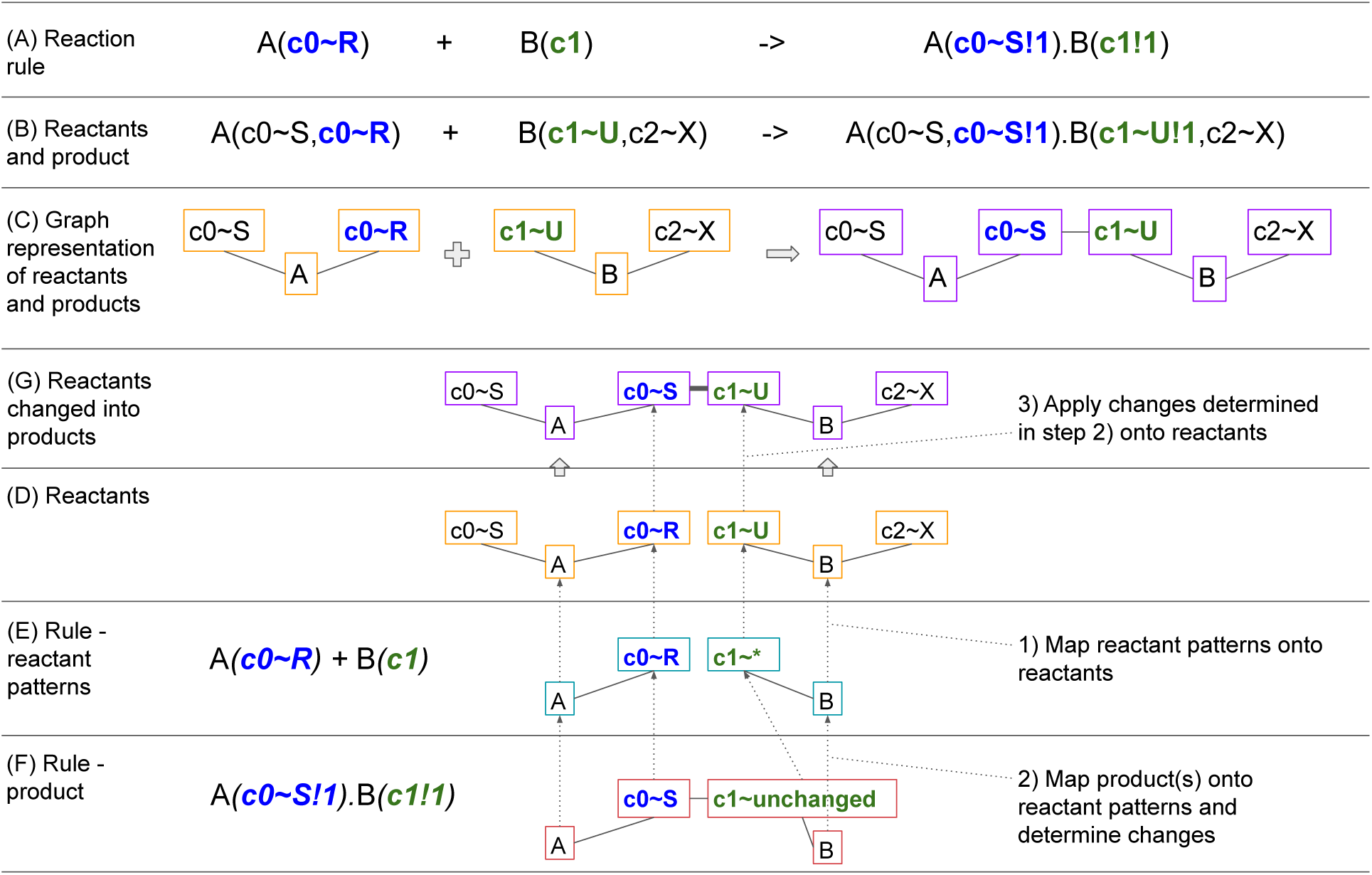
Example of a graph transformation with BNG reaction rules. In this example, reactants are defined with molecule types A(c0*∼*R*∼*S,c0*∼*R*∼*S) and B(c1*∼*U*∼*V,c2*∼*X*∼*Y) where A and B are names of the molecule types, c0 is a component of A that can be in one of the states R and S, and similarly c2 and c3 are components of B. (A) is the example reaction rule, (B) are example species reactants and products in the BNGL syntax, and (C) shows a graph representation of the rule in (B). Application of the rule is done in the following steps: 1) a mapping from each molecule and each component from reactant patterns (E) onto reactants (D) is computed (dotted arrows), if the state of a component is set in the pattern, the corresponding reactant’s component state must match. The next step 2) is to compute a mapping of the rule product pattern (F) onto reactant patterns (E). The difference between the reaction rule product pattern and the reactant patterns tells what changes need to be made to generate the product. In this example, a bond between A’s component c0 with state R and B’s component c1 is created. The state of A’s component c0 is changed to S. Once the mappings are computed, we follow the arrows leading from the reaction rule product pattern (F) to reactant patterns (E) and then to reactants (D) and 3) perform changes on the reactants resulting in the product graph (G). Each graph component of the product graph is a separate product and there is exactly one product in this example.

This graph-based approach is essential when dealing with combinatorial complexity. To model a protein that has 10 sites, in which each can be unphosphorylated, phosphorylated, or bound to another protein with ordinary differential equations (ODEs) requires 3^10^ (i.e. 59049) simultaneous ODEs [24]. For comparison, a BNGL model of the same protein will have just 6 reversible reaction rules (assuming no interaction between these 10 sites). Such a model can then be simulated using network-free simulation methods [25].

#### 2.4.1 Extension of BNGL for Volume-Surface Reactions

BNGL compartments [26] allow the definition of hierarchical volumes or surfaces where simulated molecules are located. To model a transport reaction that moves a volume molecule from one compartment through a channel (located in a membrane) into another volume compartment, one must specify the direction of this transport. We show such a reaction which implements hierarchy of compartments in Fig. 9.

**Figure 9:**
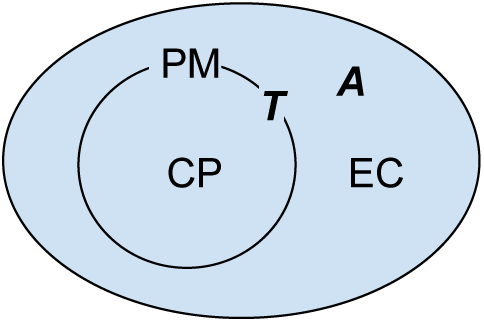
An example of compartments: EC is extracellular space, PM is the plasma membrane, and CP is cytoplasm. A is a molecule that diffuses freely in 3D space, and T is a molecule located in the plasma membrane.

In BNGL, a reaction that defines the transport of A from compartment EC into CP through transporter T is represented with the following rule:

A@EC + T@PM -> A@CP + T@PM

To model multiple instances of cells or organelles, this definition needs to be replicated with different compartments as follows:

A@EC + T@PM1 -> A@CP1 + T@PM1

A@EC + T@PM2 -> A@CP2 + T@PM2

…

MCell3 uses a general specification of orientations [4] in which the rule above is represented as:

`A’ + T’ -> A, + T’`

On the reactant side of the reaction, A’ (A followed by an apostrophe) means that molecule A hits molecule T from the “outside” (as defined below) of the compartment, and T’ means that the surface molecule T must be oriented in the membrane facing towards the outside. On the product side of the reaction, A, (A followed by a comma) means that the product A will be created on the inside of the compartment and T’ means that T will still be oriented towards the outside. Geometric objects in MCell are composed of triangles. The “outside” of a triangle is defined as the direction in which the normal vector of the triangle points. More details on molecule orientations defined in MCell3 can be found in [4].

Because the MCell3 representation of orientation is not compatible with the grammar of BNGL, and to avoid repetition of reaction rules for each compartment, we have defined an extension to BNGL that allows two special compartment classes called @IN and @OUT to be used in MCell4. With this extension reactions with compartments are then more generally defined as:

A@OUT + T -> A@IN + T

Note that only bimolecular reactions where a volume and a surface reactant interact may use the @IN or @OUT compartment classes. When the rule is used at simulation time the actual membrane compartment containing the surface reactant (T here) along with the volumetric compartment containing the volume reactant (A here) are used to correctly interpret the geometric meaning of the @IN and @OUT compartment class associated with the volume reactant. For example when this rule is applied to reactants A@EC (i.e A located in EC) and T@PM (i.e. T located in PM) at simulation time, MCell4 will first interpret the @OUT compartment class in the rule and find that compartment EC is outside of PM, and satisfies the left-hand side of the rule. Next MCell4 finds that that compartment CP is inside of PM, and finalizes the mapping of the generic compartment class @IN to the specific compartment class @CP MCell4 then inserts this specific compartment information into the rule A@OUT + T -> A@IN + T to get the runtime rule A@EC + T@PM -> A@CP + T@PM which is the same as the example rule we started with.

One more situation that we considered is how to define the orientation of the transporter in the membrane. One might need to model flippases and floppases (e.g., [27]) that change the orientation of a receptor in a membrane. In MCell3, this is handled by an orientation syntax in which a comma indicates an inward-facing orientation, and an apostrophe indicates an outward-facing orientation. In MCell4, when a molecule is created in a membrane, its orientation is always facing outwards (equivalent to T’ in the MCell3 notation). If one needs to define orientation explicitly, a component of an elementary molecule can be defined. For example one can extend the definition the molecule type T to contain a component called ‘o’ with two states called INWARDS and OUTWARDS. The rule defined for a specific state of the transporter will then be:

A@OUT + T(o∼OUTWARDS) -> A@IN + T(o∼OUTWARDS)

To flip the orientation of T, a standard BNGL rule F + T(o*∼*OUTWARDS) -> F + T(o*∼*INWARDS) can be defined; Here, F is a surface molecule flippase.

To summarize, we introduced an extension to BNGL in which compartment classes @IN and @OUT are used to define general volume+surface molecule reaction rules that can be applied to any specific compartments at simulation time.

#### 2.4.2 Units and Interoperability between MCell4 and BioNetGen

Usage of the BioNetGen language offers an excellent interchange format. Model definitions in BNGL can be executed by MCell, and BioNetGen itself implements various simulation approaches such as ODE, SSA, PLA, and NFSim. BioNetGen does not have pre-described units so that the user is free to use any unit system they deem suitable and that is compatible with the underlying algorithms. To facilitate model interchange, we define a set of units to be used when BNGL models are implemented in MCell4 and when the model is exported for use within BioNetGen as shown in Table 1.

**Table 1:** Units used in MCell and suggested units for BioNetGen. Unit N represents the number of molecules and M is molar concentration. BioNetGen interprets membranes (2D compartments) as thin volumes of thickness 10 nm. NFSim in BioNetGen does not fully support compartmental BNGL yet and the volume of the compartment must be incorporated into the rate units of the reactions occurring in that compartment, therefore NFSim’s bimolecular reaction rate unit does not contain a volumetric component. Additional units in MCell include: length in µm and diffusion constants in cm^2^ s^−1^.

| Simulation tool and mode of usage | Volume-volume or volume-surface bimolecular reaction rate | Surface-surface bimolecular reaction rate | Unimolecular reaction rate | Compartment volume | Seed species (initial molecule release) value |
| --- | --- | --- | --- | --- | --- |
| MCell4 with default units | $M^{-1} s^{-1}$ | $\mu m^2 N^{-1} s^{-1}$ | $s^{-1}$ | $\mu m^3$ | N |
| MCell with BNG units; BioNetGen ODE, SSA, PLA | $\mu m^3 N^{-1} s^{-1}$ | $\mu m^3 N^{-1} s^{-1}$ | $s^{-1}$ | $\mu m^3$ | N |
| BioNetGen NFSim | $N^{-1} s^{-1}$ | $N^{-1} s^{-1}$ | $s^{-1}$ | ignored | N |

An MCell4 model is typically implemented as a combination of Python and BNGL code. Although the approach that we recommended is to capture all the reaction rules and initial molecule states in BNGL, it may sometimes be beneficial to use Python code for these definitions (e.g., to generate reaction networks programmatically). There are also aspects of spatial models that cannot be captured by BNGL. To simplify model validation, MCell4 provides an automated means to export a model that has been implemented as a combination of Python and BNGL into pure BNGL. Since not all features (especially spatial distributions) of an MCell4 model can be mapped to pure BNGL, a best-effort approach is used during this export. All model features that can be translated into BNGL are exported and error messages are printed identifying the model aspects that have no equivalent in BNGL. If the exported model includes all essential model aspects it can be used for cross-validation and comparison of results between the MCell4 and BioNetGen simulations. Verifying results with multiple tools can reveal errors in the model or in the simulation tools. Therefore, such validation is a recommended step in development of an MCell model.

#### 2.4.3 Example of an MCell4 Model with BioNetGen Specification

To demonstrate the support for BNGL in MCell4, we show a simple example (Fig. 11) that imports (i.e. loads) information on species and reaction rules, molecule releases, and information about compartment from a BNGL file (Fig. 10).

**Figure 10:**
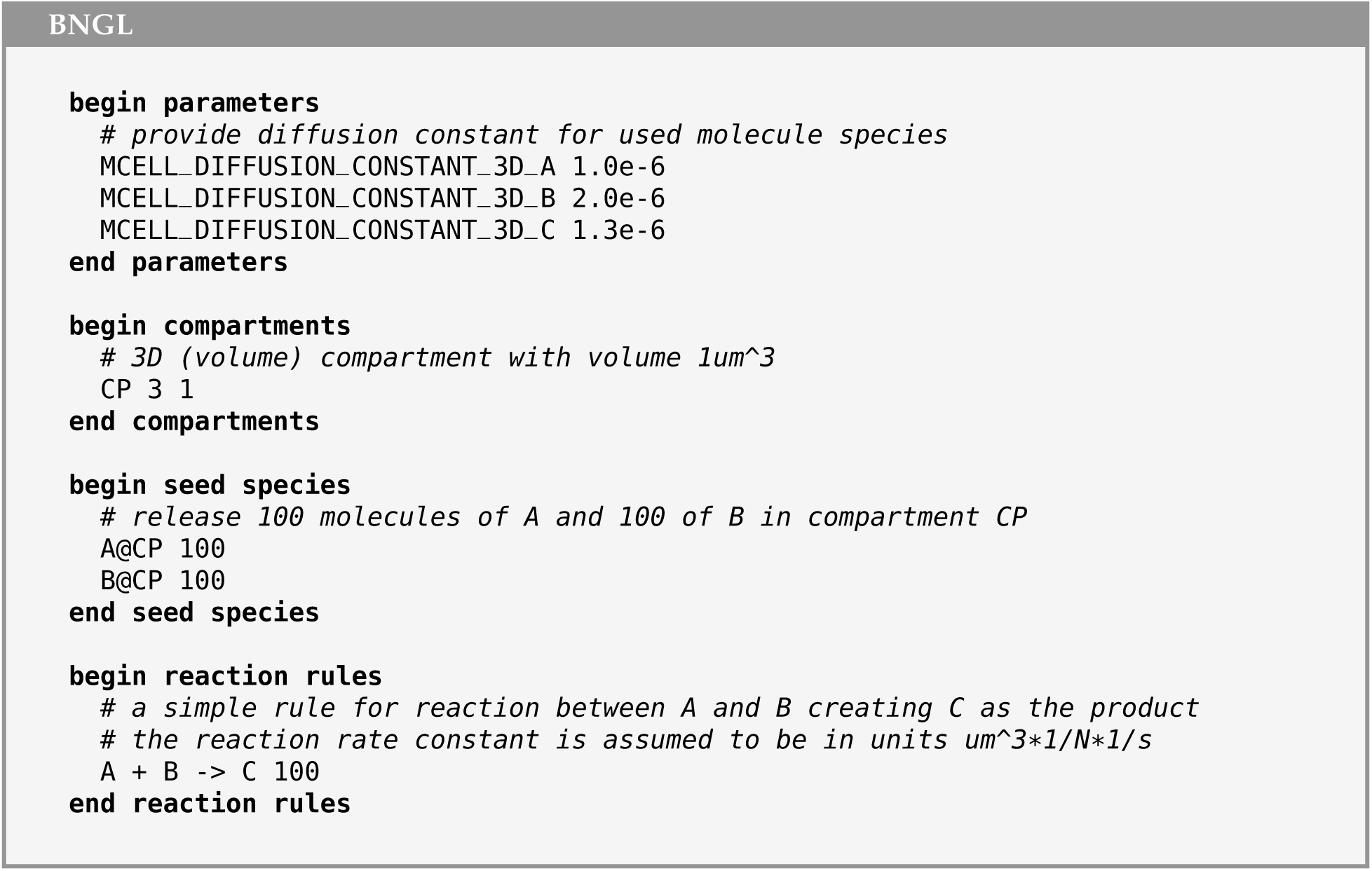
BNGL file that defines a compartment CP, and instantiates release of 100 molecules of A and 100 molecules of B into it. It then implements a reaction rule in which A and B react to form the product C.

**Figure 11:**
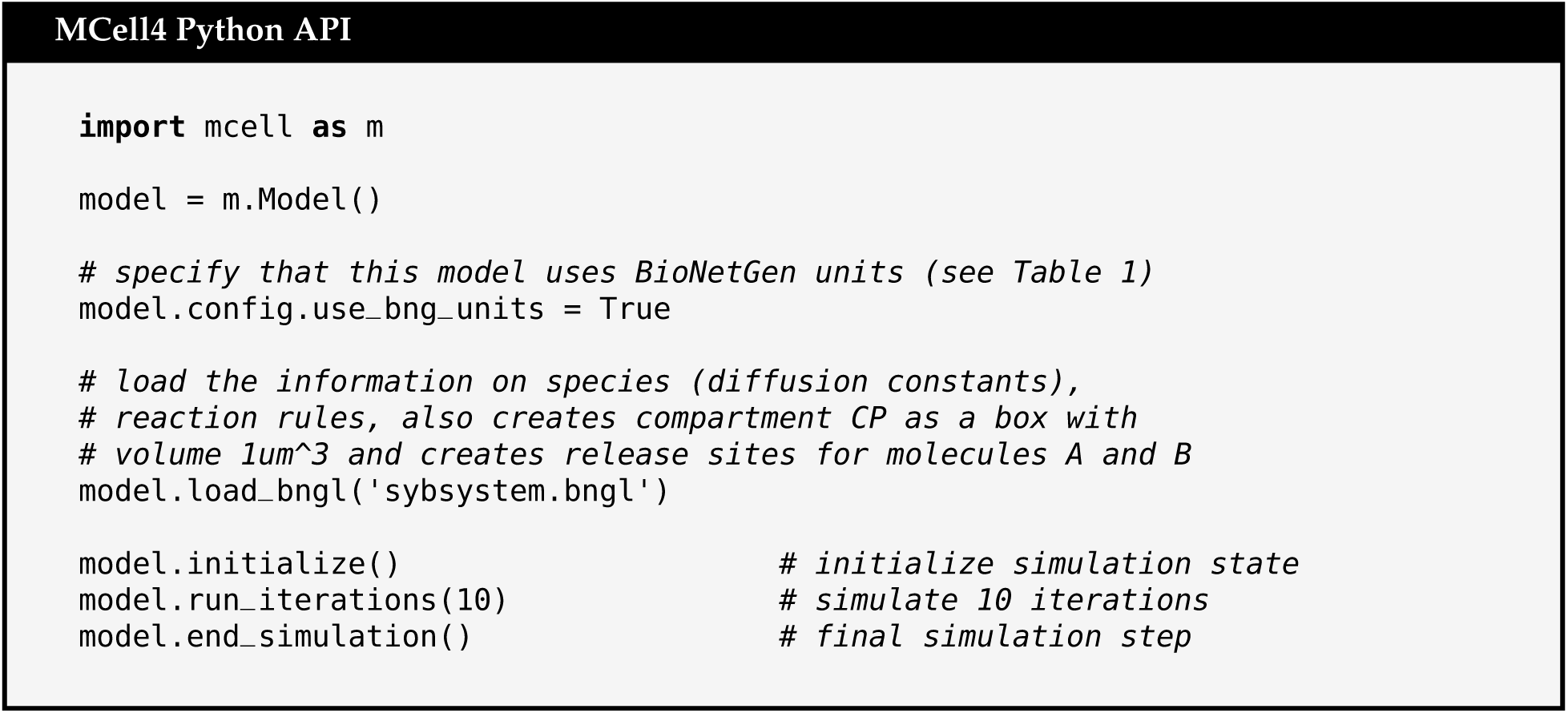
Python code for an MCell4 model that will implement loading of the BNGL file shown in Fig.10 (referenced as subsystem.bngl). In this example the entire BNGL file is read. It is also possible to load only specific parts of the BNGL file, for example only reaction rules or only compartment and molecule release information. It is also possible to replace BNGL compartments with actual 3D geometry.

Note that the file in Fig. 10 is a standard BNGL file that can be used directly by other tools such as BioNetGen so that no extra conversion steps are needed for the BNGL file to be used elsewhere. This permits fast validation of a reaction network with BioNetGen’s ODE or other solvers. The model can be checked against the spatial simulation results in MCell4 without the need to have multiple representations of the same model.

## 3 Results

### 3.1 Testing & Validation

We performed extensive testing and validation to ensure the accuracy of results generated by MCell4. We compared results from the previous versions, MCell3 [4] and MCell3-R [12], which were themselves extensively tested prior to their release. One can obtain byte for byte identical results with MCell3/MCell3-R and MCell4 by using specific options during compilation. These options ensure that the molecules are simulated in the same order and with the same stream of random numbers in MCell3/MCell3-R and MCell4. We have created a test suite (included in the MCell source code repository) containing more than 350 validation tests that verify correct results in MCell3 and MCell3-R tests. We obtain byte for byte identical results of these tests between MCell3, MCell3-R and MCell4. Simulation results were also validated against results with a BioNetGen ODE solver [6] and with NFSim [14] by running equivalent models in MCell4 and in BioNetGen, running with up to 1024 different random seeds. The diffusion constants in MCell4 were set to a high value to emulate a well-mixed solution. We then compared the shape of the time course of the averaged counts (and variance of counts) of molecules of a given species. Some tests cases have an analytic solution. The results of all tests agreed well between simulators, and with analytic solutions, and were always within the measured variance in all cases. More than 45 of such tests are included in the MCell4 test suite [28]. Some of these tests are referenced as examples in MCell4’s API reference manual [29].

#### 3.1.1 SNARE Complex

We implemented a model of the SNARE complex, a cooperative dual Ca^2+^ sensor model for neurotransmitter release [30], as an example of an MCell4 model containing a BioNetGen specification. The model includes the binding of up to five calcium ions to the sensor and synchronous or asynchronous modes of release of neurotransmitters. An adapted version of this model was previously implemented in an older version of MCell [31]. The model is composed of SNARES with 18 state variables, calcium ions and 63 reactions. There are different possible implementations of the model in BNGL. The one presented here is compatible with MCell4, and allows simulation of the model in BioNetGen and MCell4 without modifying the code. It consists of three molecules types and ten reaction rules (Fig. 12). The snare complex (represented as **snare**) is an elementary molecule that has eight components: five **s**, that represent the binding site for calcium molecules in the synchronous sensor; two **a** components that represent the binding sites for calcium in the asynchronous sensor; and one component called **dv** with two states (*∼* 0 *∼* 1), that represents docking of a vesicle to the snare complex (*∼* 1) or its absence (*∼* 0). Calcium ions (Ca^2+^) can bind and unbind to the complex. The release of neurotransmitters is tracked via a dummy molecule type called V_release(), which captures the timing of the release but does not actually release molecules of neurotransmitter (see the next section for an implementation of the release in MCell4). Fig 13A shows code implementing the states of the model, and the synchronous and asynchronous release. Assuming well-mixed conditions, a large volume containing the surface complexes, a large number of complexes and a constant calcium concentration, the results obtained with BioNetGen ODE simulations and the spatial model in MCell4 give qualitatively similar results (Fig 13B). The source code for this example can be found in [32].

**Figure 12:**
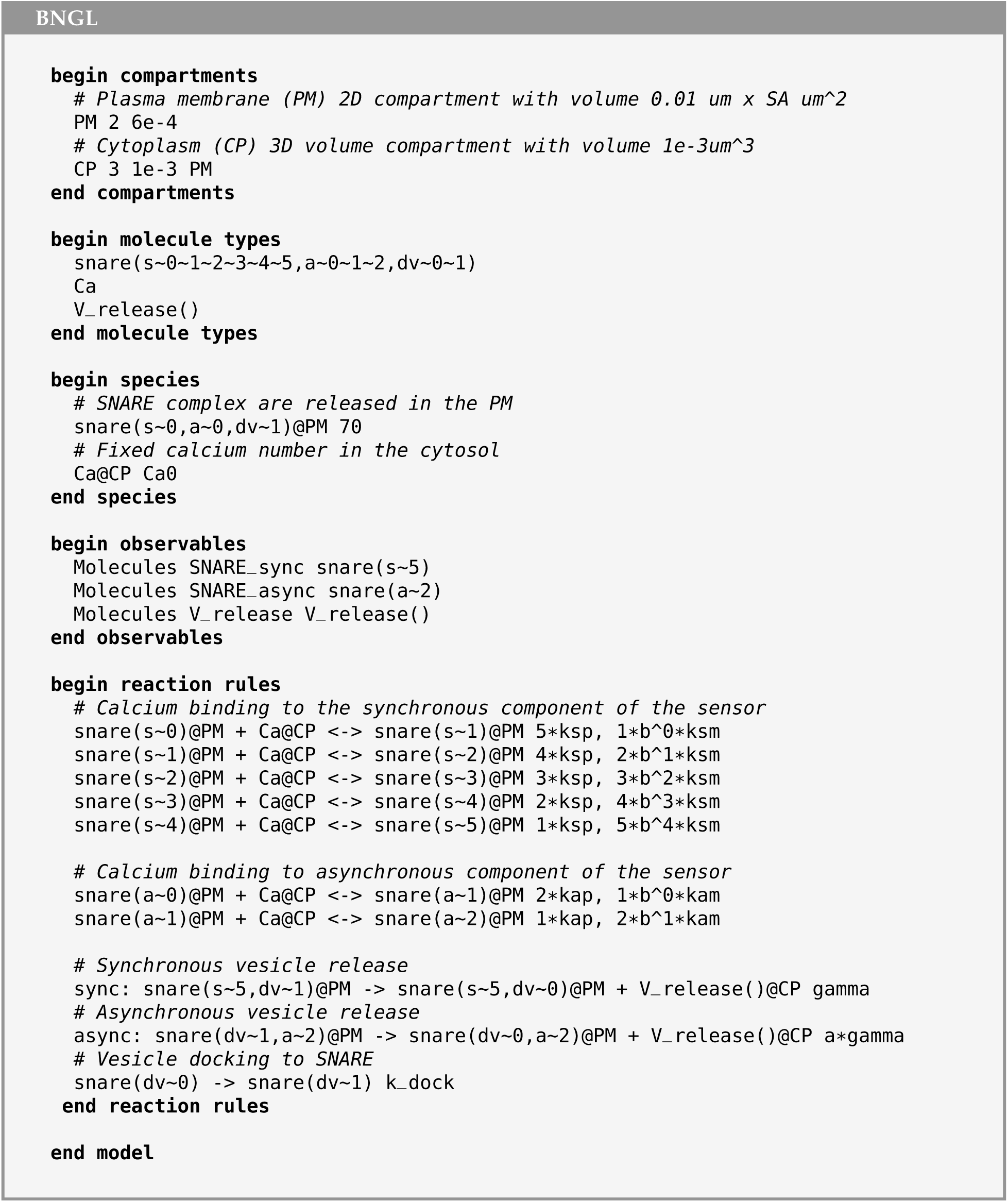
Compartmental BNGL implementation of the SNARE complex model. One 3D compartment, cytosol (CP), and its associated plasma membrane (PM) is defined. Molecule types are defined, and their released sites are specified: SNARE molecules are released into the PM, and Calcium ions into the Cytosol. This code is followed by specification of the observables, and the reaction rules governing the interactions.

**Figure 13:**
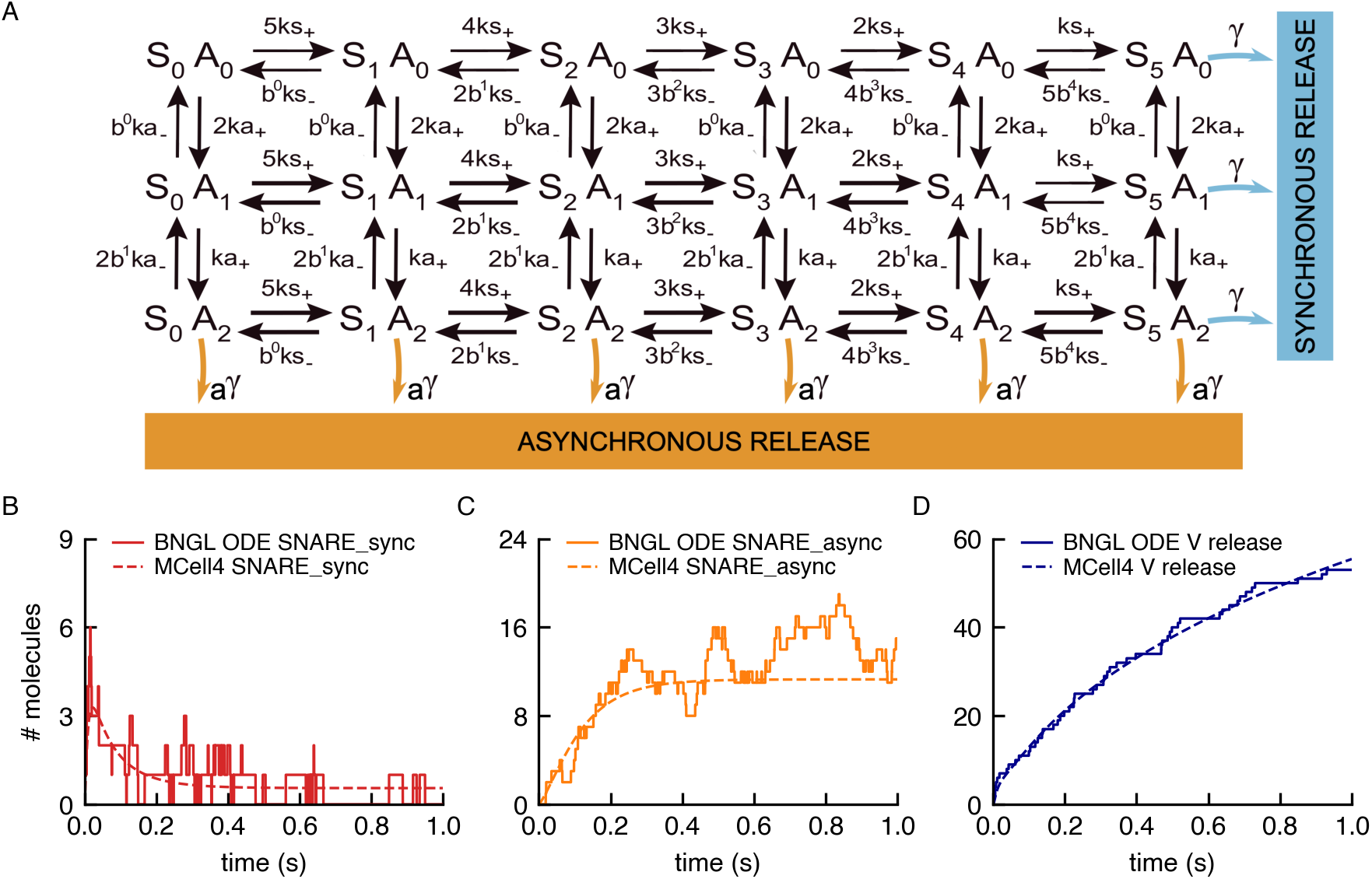
(A) Schematic diagram of the state variables of the SNARE complex model. It consists of 18 states, S and A represent the synchronous and asynchronous components of the complex, which can be in five and two different states respectively (B-D) Results of independent simulations of the model with ODEs in BioNetGen (dashed lines) and in MCell4 (solid lines).

#### 3.1.2 Event-Driven Release of Neurotransmitter by the SNARE Complex

To release neurotransmitter in an event-driven manner at the times captured by observing V_release() in the SNARE example above, we employ a new feature in MCell4: callbacks. One of the most powerful new features of MCell4 is the ability to implement python code to be executed (i.e. called) each time a user-specified reaction or wall collision event occurs during a simulation; thus, the term “callback”. In the MCell4 Python API the code to be executed when “called” by the event, is written as a function and this function is referred to as a “callback function”.

In this case we created a callback function that will release a given number of neurotransmitter molecules, at the time the synchronous or asynchronous reactions occur. We localize the release at the position of the individual SNARE complex that triggers the release. Here we briefly describe how this is accomplished. The full details and Python source code of the working MCell4 model can be found at https://github.com/mcellteam/article_mcell4_1/tree/master/snare_complex/snare_w_callback.

There are two types of callback functions supported in the MCell4 Python API, “reaction callback functions” and “wall hit callback functions”. In the SNARE complex example we have created a reaction callback function that will be called upon the stochastic occurrence of the “sync” or “async” reactions specified in Fig. 12. We name this reaction callback function “release_event_callback” and associate it with the reactions using the “register_reaction_callback()” command provided in the API. During simulation of the model, whenever a sync or async reaction occurs, the MCell4 physics kernel will execute “release_event_callback()”. To specify which species of neurotransmitter to release, how much, and where the register_reaction_callback() command allows additional metadata (called “context”) to be passed to the callback function. In this example we created a Python class called “ReleaseEventCallbackContext” which contains the name of the species and number to be released, as well as the relative release location. Release_event_callback() can then make use of this context to perform the desired operations. See file “customization.py” in the working model for complete details.

#### 3.1.3 CaMKII Model with Large Reaction Network

To demonstrate results for a system with a large reaction networks, we use a model of a CaMKII dodecamer which is an extension of a model described in [33].

The CaMKII dodecamer (a “protein complex”) is composed of two CaMKII hexameric rings stacked on top of each other. Each CaMKII monomer with its calmodulin (CaM) binding site can be in one of 18 states. Then the total number of states possible for a CaMKII dodecamer CaMKII is then 18^12^ */*12 *≈* 10^12^ (the division by 12 is to remove symmetric states). This is an example of the combinatorial complexity mentioned in section 2.4 for which it is simply not feasible to expand all the reaction rules and generate the entire reaction network to be stored in memory, and thus a network-free approach is necessary. Fig. 14 shows the results of validation of this model against BioNetGen/NFSim, MCell3R, and MCell4.

**Figure 14:**
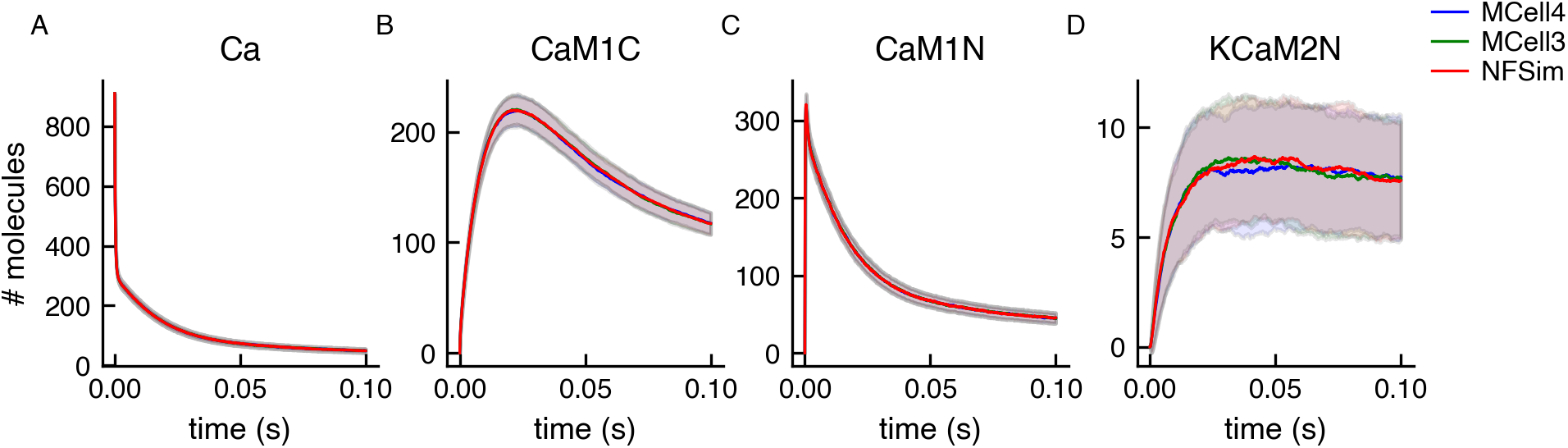
Validation of MCell4 simulation against BioNetGen/NFSim and MCell3R using a CaMKII model. The input BNGL model for NFSim was obtained by automatic BNGL export of BNGL from the MCell4 model. The simulation ran for 100000 iterations (0.1 s). Lines in the graphs are averages from 256 runs with different random seeds, and bands represent one standard deviation. Molecules in MCell3R and MCell4 use diffusion constant of 10*^−^*^3^ *cm*^2^ */s* to emulate a well-mixed solution (the usual value is around 10*^−^*^6^ *cm*^2^ */s*). The names of the observed species are indicated in the graph titles: CaM1C is CaM(C*∼*1, N*∼*0, camkii); CaM1N is CaM(C*∼*0, N*∼*1, camkii); KCaM2N is CaMKII(T286*∼*U, cam!1).CaM(C*∼*0, N*∼*2, camkii!1). The simulation was initiated far from equilibrium; therefore there was an initial jump in the molecule numbers. The molecule names are explained in [33].

We also present an extension of the aforementioned model [33], in which we can now observe the effects of the geometry of the compartment on the simulation results by modeling in MCell4. Figure 15 shows three different variations of the model. The first variation distributes the molecules homogeneously in the compartment (equivalent to the well-mixed versioned published previously, Figure 15 A). Two additional variations include a small subcompartment, located near the top of the larger compartment, that is not transparent to diffusion of CaMKII and CaM molecules. In the first variation all the molecules are homogeneously distributed throughout the compartment, but the the CaMKII and CaM molecules in the subcompartment do not mix with the rest of the compartment (Figure 15 B). In the second variation, half of the CaMKII molecules are placed in the subcompartment and the other half in the remainder of the compartment, while CaM is still distributed homogeneously throughout the entire volume (Figure 15C).

**Figure 15:**
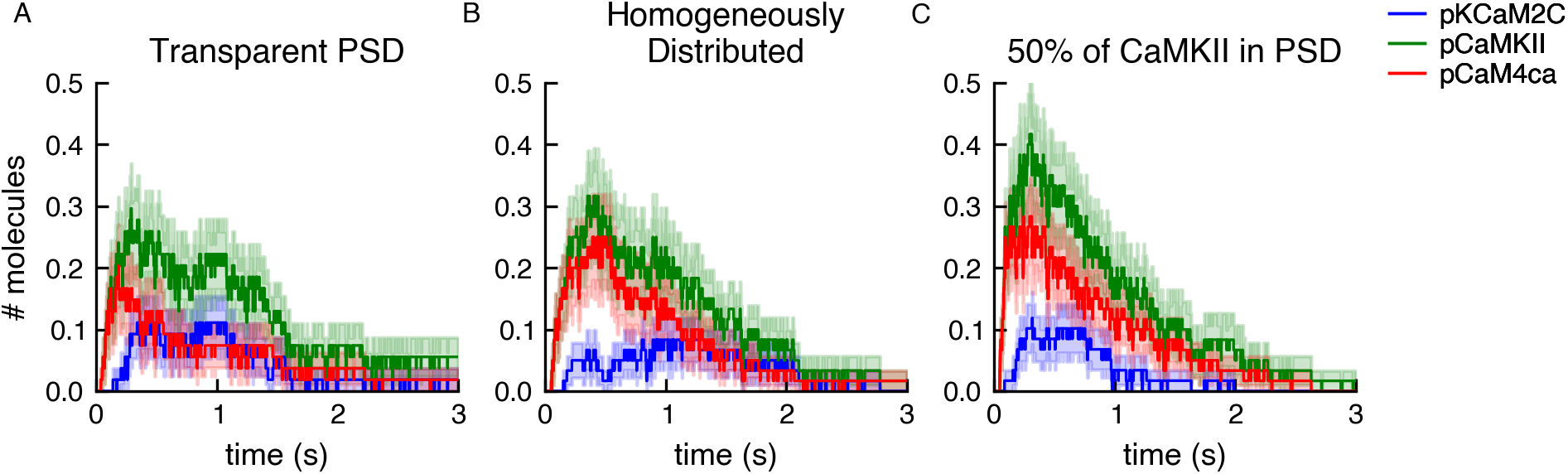
The effect on CaMKII phosphorylation of trapping CaMKII and CaM inside a subcompartment named PSD. Three different conditions where simulated. (A) All molecules are homogeneously distributed throughout the entire compartment. (B) A small subcompartment, termed PSD, which is reflective to CaMKII and CaM, but is transparent to calcium ions and PP1, is added near the top of the larger compartment. All the molecules are homogeneously distributed throughout both compartments. (C) The subcompartment is reflective to CaMKII and CaM and 50% of the CaMKII molecules are trapped inside the subcompartment, and the rest of the molecules are distributed homogeneously throughout the remainder of the larger compartment. The plots show an average of 60 runs, lighter shaded bands represent standard error of the mean.

We sought to observe the effect of these three conditions on CaMKII phosphorylation as a result of Ca^2+^ influx into the compartment. In all three conditions a Ca^2+^ influx is simulated from a single point source located in the center of the top face of the large compartment. As in [33] the Ca^2+^ influx was such that at the peak the free calcium concentration was *∼* 10*µ M*, and it returned to near the steady state level within 100 ms. These spatial differences have a small but significant effect on CaMKII phosphorylation levels in response to the Ca^2+^ influx. These differences would have been impossible to investigate without the combination of the network-free simulations and the diffusion in space implemented in MCell4.

#### 3.1.4 Volume-Surface and Surface-Surface Reactions: Membrane Localization Model

We used a membrane localization model from [34] (section 2A) to validate volume-surface and surface-surface reactions.

The model analyzes how membrane localization stabilizes protein-protein interactions. A pair of protein binding partners A and B are localized to the membrane surface by binding a lipid molecule M. This binding to the membrane constrains the space in which the molecules diffuse and thus promotes complex formation. The model is created within a box of dimensions 0.47 *×* 0.47 *×* 5*µm*^3^. Surface molecules M are released on one of the smaller sides of the box. The 4 edges of this side are set to be reflective, so the surface molecules cannot diffuse onto the other sides.

MCell subdivides the surface areas of geometric objects into small tiles. A maximum of one molecule can occupy one tile at a time - this tiling simulates volume exclusion for surface molecules. A parameter named “surface grid density” sets the density (and size) of the tiles and the thus the maximum packing density of surface molecules. The initial density of surface molecules in this model is 17000 *molecules/µm*^2^, and we set the surface grid density to 40000 *tiles/µm*^2^ giving an occupied area fraction of 42.5%. (SAY MORE ABOUT RESULTS OF VALIDATION TESTS IN FIG 16. DEFINE NERDDS. ADD SMOLDYN RESULT TO FIG 16)

**Figure 16:**
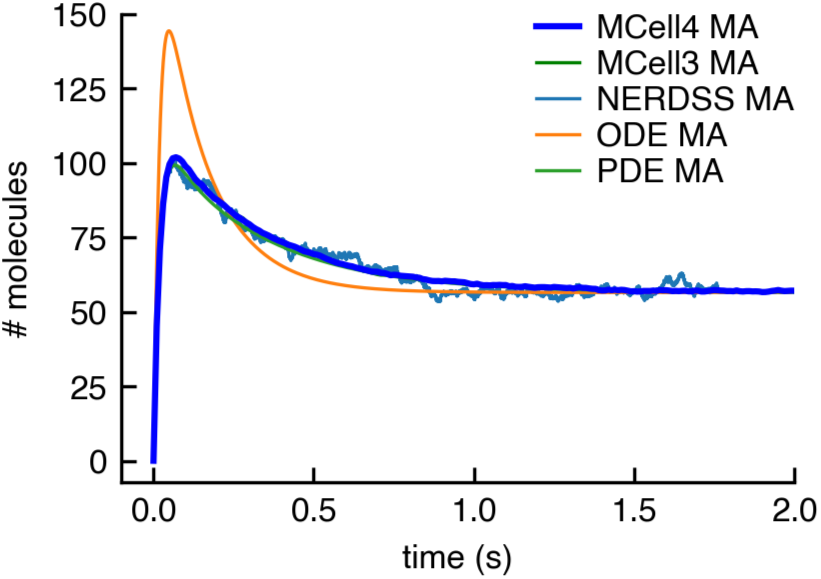
Simulation results for the membrane localization model. The plot shows copy numbers of a surface molecule MA (surface molecule M with a bound volume molecule A). MCell4 and MCell3 results show a good match with the NERDDS simulator (NERDDS results are from from [34], the data ended at time 1.75 s). The results computed with ODE and PDE solutions produced by VCell reach the same equilibrium (VCell results are from from [34] simulated with VCell 7.2.0.39). (ADD SMOLDYN RESULTS) MCell3 and MCell4 results are an average of 512 runs with different random seeds.

#### 3.1.5 Stochastic Fluctuations in a System with Multiple Steady States: Autophosphorylation

Another validation model from [34] (section 2B) shows stochastic fluctuations in a system with multiple steady states. A deterministic ODE solution does not show these multiple steady states and almost immediately stabilizes in one of them. In Fig. 17 we show the output of an MCell4 simulation and a simulation of the same model simulated in NFSim using the BNGL exported from the MCell4 model (more details on BNGL export are in 2.4.2). We also illustrate the steady states reached with ODE solutions.

**Figure 17:**
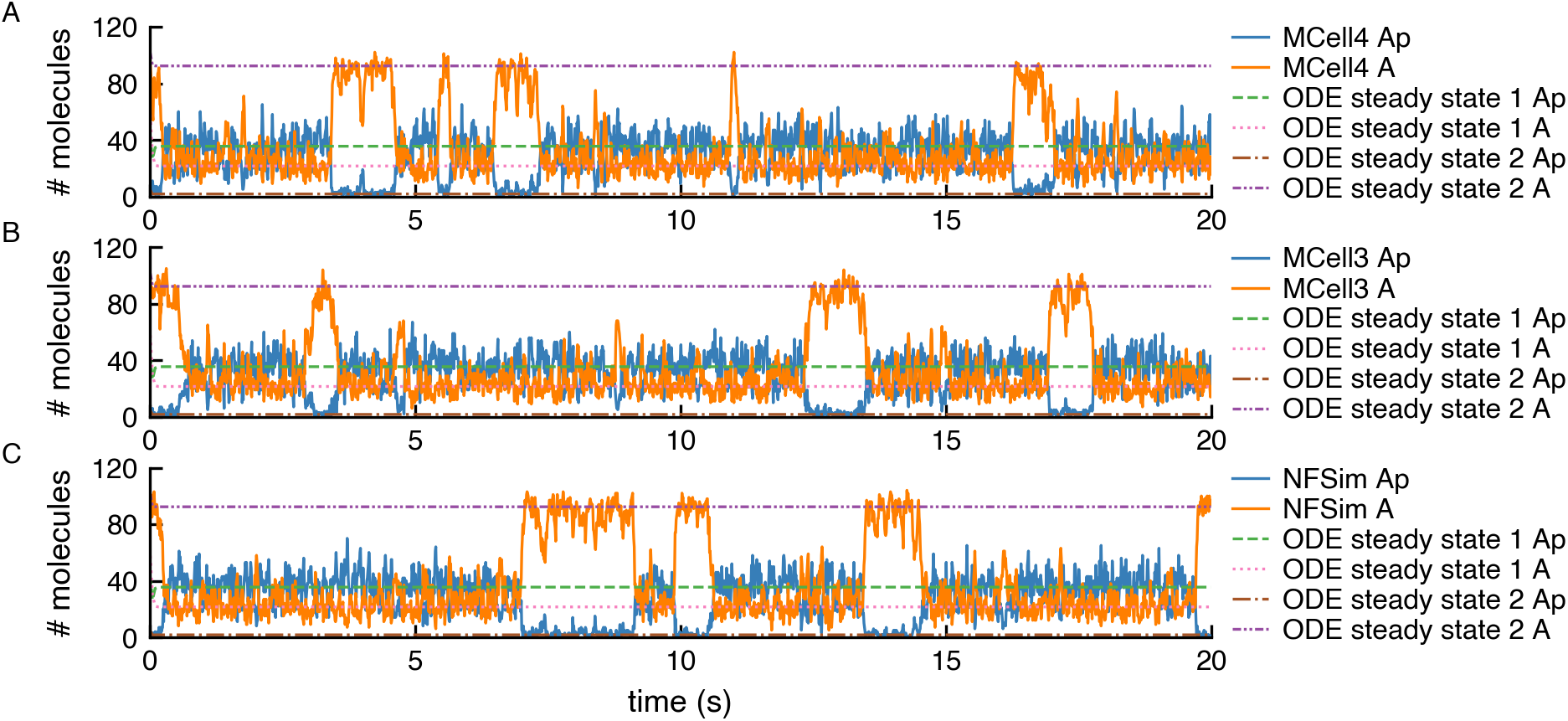
An example of a stochastic simulation of a system that exhibits switching between multiple steady states. Copy numbers of unphosphorylated kinase A and its phosphorylated variant Ap are shown for a single simulation run in MCell4, MCell3, and NFSim. The NFSim model was obtained by automatically exporting the MCell4 model into BNGL. The graphs also show solutions obtained with a deterministic ODE model for which data from [34] were used. The results demonstrate that the MCell results correctly reach one of the stable steady states shown in the ODE results. The simulation stays in such a state, and then due to stochastic behavior, a switch another steady state occurs.

### 3.2 Performance

With relatively small reaction networks (less than 100 or so reactions), the performance of MCell4 is similar to MCell3 as shown in Fig. 18 (A). MCell3 is already highly optimized. MCell3 contains optimization of cache performance that speeds up models with large geometries; this optimization is not present in MCell4. Thus MCell3 is faster for large models such as models created in neuropil reconstructions containing on the order of 4 million triangles defining their geometry. The situation is different when comparing MCell4 and MCell3-R with models that use large BNGL-defined reaction networks (18B). MCell3-R uses the NFSim library to compute reaction products for BNG reactions. With large reaction networks containing as many as 10^10^ reactions or more, MCell3-R stores all the reactions that occur during run-time in memory and thus gradually slows down. We have not been able to implement reaction cache cleanup in MCell3-R. MCell4 with the BNG library keeps track of the number of molecules of each species in the system during simulation and periodically removes from the cache reactions and species that are not used. This facilitates simulation of complex reaction networks with a potentially infinite number of species and reactions without excessive impact on memory usage and performance.

**Figure 18:**
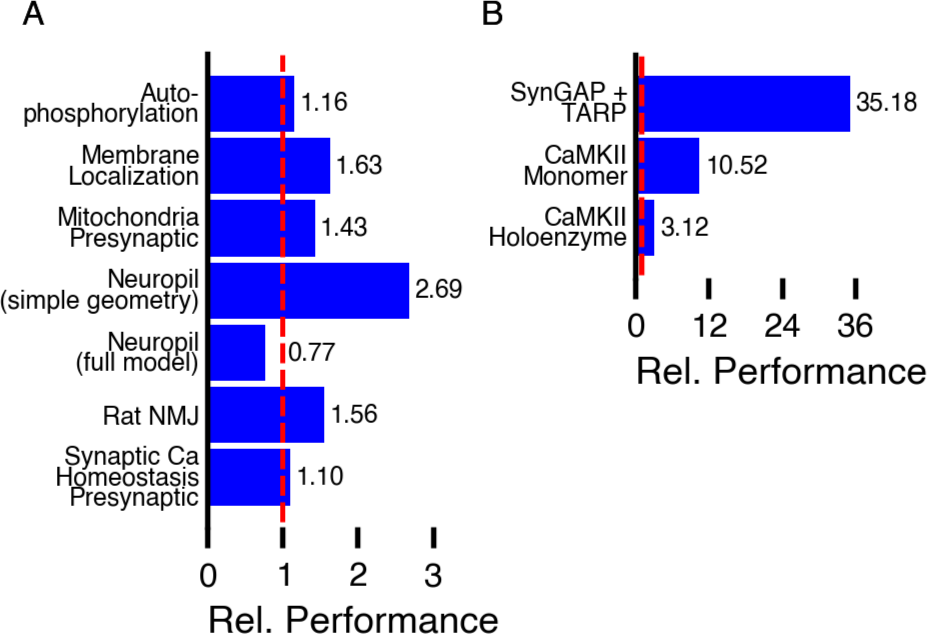
For selected benchmarks, we measured elapsed time for how long the simulation ran starting from the second iteration (after all initializations) and ending when the simulation finished. Time was measured on AMD Ryzen 9 3900X@3.8GHz. Both MCell3 and MCell4 use a single execution thread. Relative performance shown in the graphs is computed as time for MCell3 or MCell3-R divided by time for MCell4. The sources of the models are as follows: Presynaptic Ca Homeostasis [31]; Rat Neuromuscular Junction [2] model with updated geometry (shown in Fig 1), Neuropil [5]; Mitochondrion Model [35]; Membrane Localization [34]; Autophosphorylation [34]; CaMKII Monomers [33]; CaMKII Holoenzyme [33]; SynGAP with TARP (not yet published).

### 3.3 Hybrid Simulation Example

MCell4’s Python API supports interaction with an MCell4 simulation while it is running. Here we show a model in which the progression over time of one molecular species is encoded in Python code with a differential equation and the remaining species are encoded in MCell4 as particles behaving stochastically. As a basis for this demonstration we used a model of a circadian clock published in article [34], originally based on article [36].

The model simulates the behavior of an activator protein A and repressor protein R that are produced from mRNA transcribed from a single copy of a gene (one for each protein). Coupling of A and R expression is driven by positive feedback of the activator A, which binds to each gene’s promoters to enhance transcription. Protein R also binds to A to degrade it. All other proteins and mRNA are degraded spontaneously at a constant rate.

Compared to the original model in [36], authors of [34] increased the reaction rates in the model from hours to seconds by multiplying the reaction rates by 3600. Because the purpose of this example is to demonstrate a hybrid model in MCell4 and its validation, which requires many runs, we made another change to accelerate the simulation; we reduced the simulation volume by a factor of 268 to 0.25 µm which increased the rate of bimolecular reactions. We also increased the unimolecular reaction rates by the same factor.

In the hybrid model, protein R is simulated as a changing concentration, under well-mixed conditions, whose concentration value is updated by finite difference expressions. The other species are simulated as particles. In the base MCell4 model, there are 4 reactions that consume or produce R (Fig. 19). We replaced two of these with reactions that do not model R as a particle and the remaining two reactions were replaced with finite difference expressions Fig. 20). The hybrid coupling of the finite difference calculations with MCell4’s particle-based calculations is shown in the pseudo-code representing the main simulation loop in Fig. 21.

**Figure 19:**
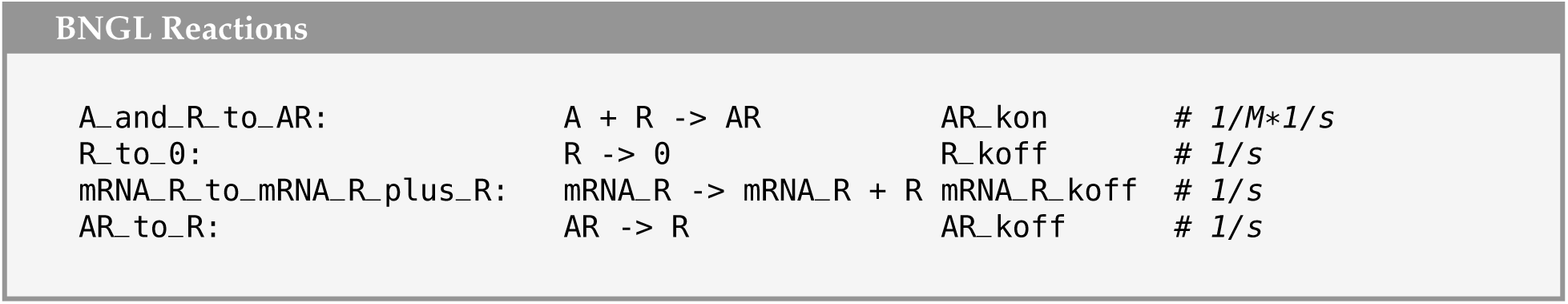
Reaction rules affecting protein R in the particle-only model.

**Figure 20:**
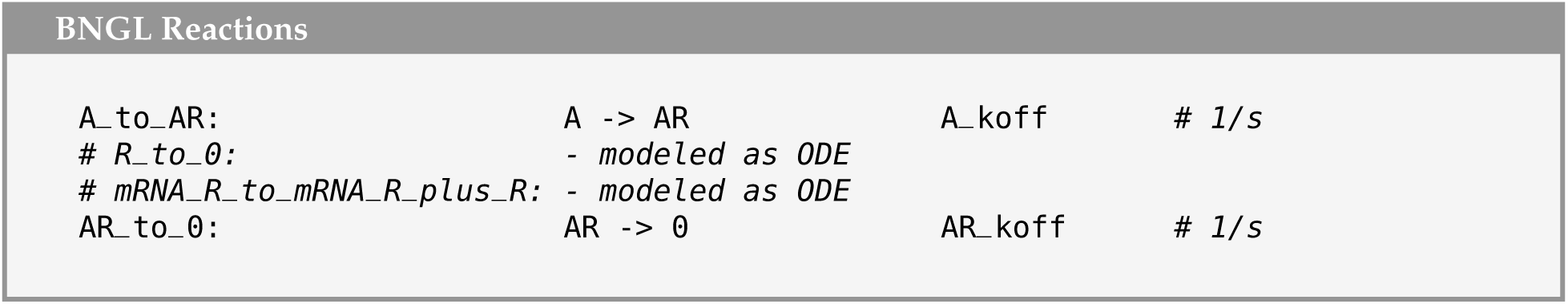
Reaction rules affecting protein R in the hybrid model.

**Figure 21:**
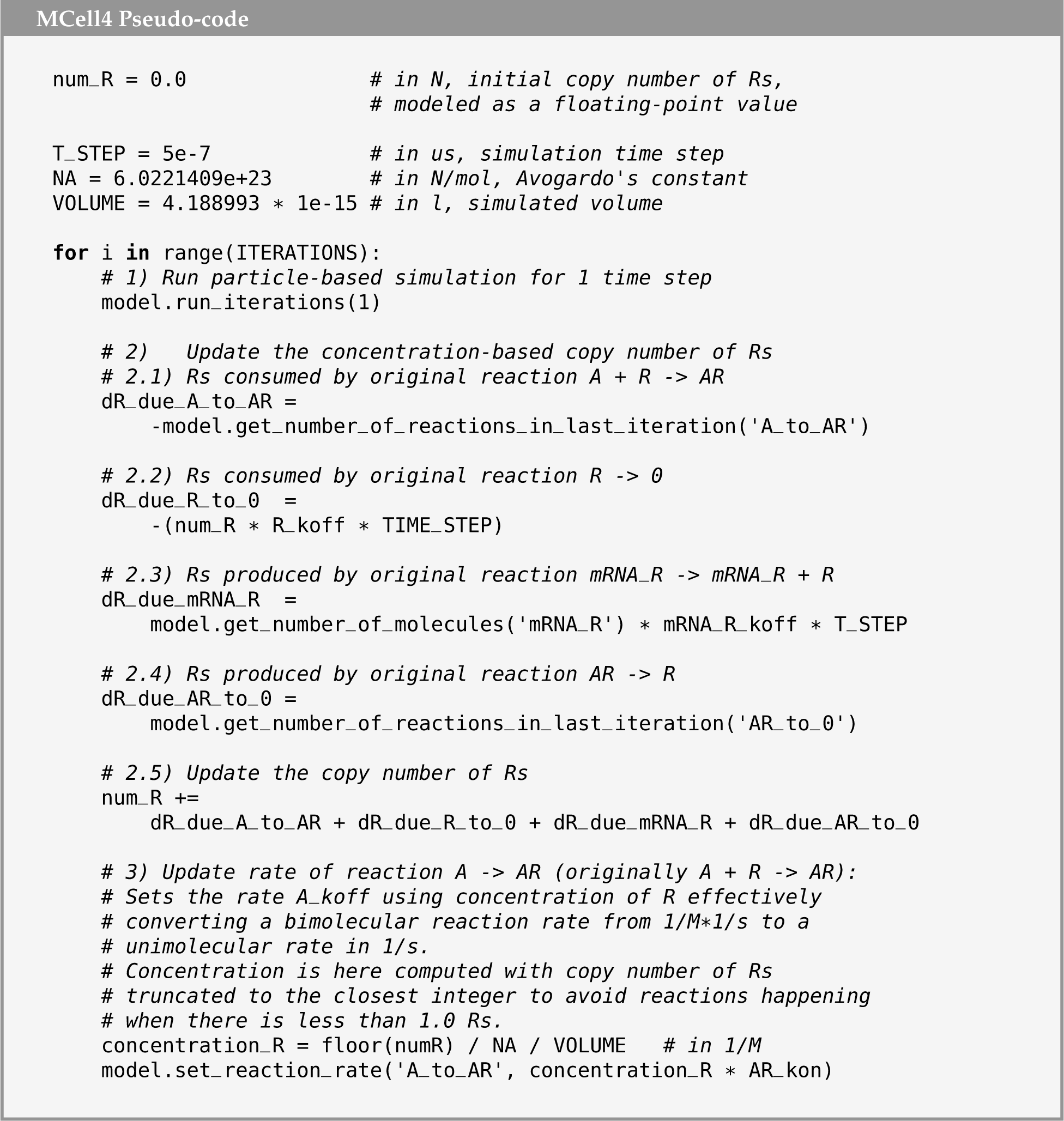
Pseudo-code of the main simulation loop that: 1) runs an iteration of the particle-based simulation, 2) updates the copy number of R based on the current MCell state, and 3) updates the rate of reaction A -> AR that was originally a bimolecular reaction A + R -> AR. N is a unit representing the copy number. This pseudo-code was adapted to show the actual computations in a more comprehensible way. The runnable MCell4 Python code is available in the GitHub repository accompanying this article [32].

To validate that the results of the hybrid variant of the model are correct, we ran 1024 instances of stochastic simulations with different initial random seeds. We also compared the effect of two different diffusion constant values when using MCell. Results that showing the average oscillation frequencies are shown in Fig. 22 and the copy numbers of molecules A and R in Fig. 23.

**Figure 22:**
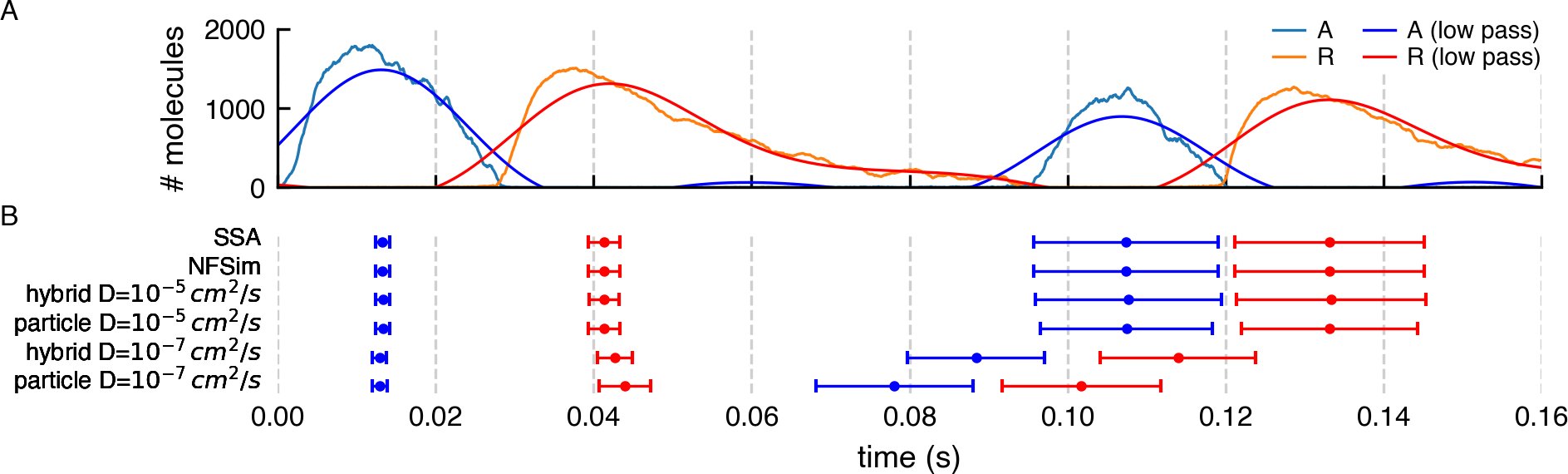
(A) Result of a stochastic simulation of a circadian clock model with NFSim. Copy numbers of molecules A and R show periodic oscillation. A low pass frequency filter was used to smooth the values of A and R. The reason for the smoothing was to get a numerical value related to the actual peak The peaks from low-pass filtered data do not represent actual average peaks but can be used as a proxy to obtain the time of a peak for comparison with other simulation methods. (B) The error bars capture the mean and standard deviation of the low pass filtered peak times for different variants of the model and simulation algorithms. Each of the variants was run 1024 times. It is evident that the SSA, the NFSim, and the MCell model variants with a fast diffusion constant, *D* = 10*^−^*^5^ *cm*^2^ */s*, give essentially the same results. The hybrid MCell model with the slower diffusion constant, *D* = 10*^−^*^7^ *cm*^2^ */s*, shows faster oscillation than the non-spatial models run with SSA and NFSim, and the MCell4 variants with faster diffusion. The pure particle-based MCell4 model with *D* = 10*^−^*^7^ *cm*^2^ */s* shows the fastest oscillations.

**Figure 23:**
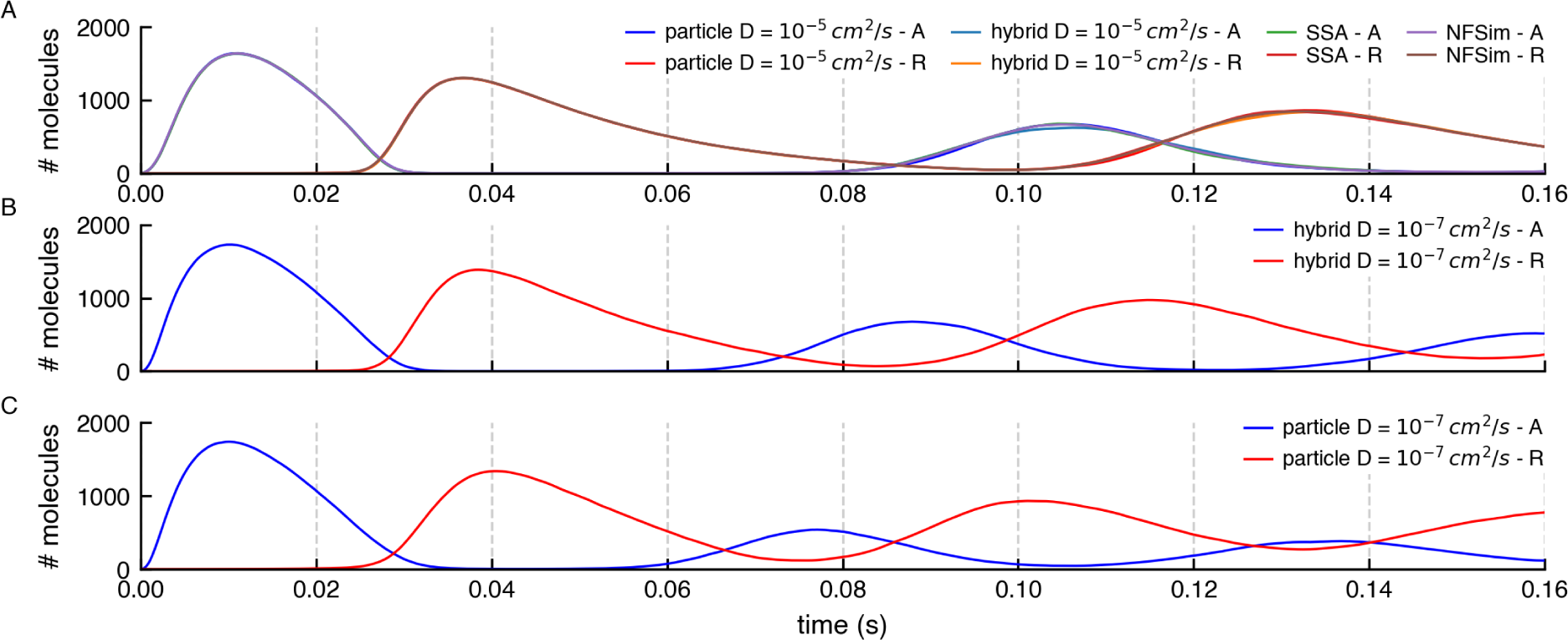
Comparison of copy numbers of A and R during simulations by different methods. (A) The average copy numbers for A and R proteins from 1024 runs in NFSim, SSA, and MCell4 with a fast diffusion constant match. To get an even better match,would require more than 1024 runs because stochastic molecular simulations show high variability when the copy number of some of the species is low which is the case here for both A and R. (B) and (C) Average copy numbers for MCell4 simulations with a slow diffusion constant. These are shown as separate plots to highlight the effect of slow diffusion on spatial simulation results.

When using a fast diffusion constant of 10*^−^*^7^ *cm*^2^ */s* for all molecules, all simulation approaches produce essentially the same results. A significant advantage of using hybrid modeling is often that the hybrid model runs much faster, as in this specific example, in which the simulation speed of the MCell4 hybrid model is 4x faster. This is because: 1) The time step can be set to 5x longer because there is no need to model explicitly the diffusion of particle-based molecules for the fastest reactions. Note that the time step when all molecules are modeled as particles must be 10*^−^*^7^ *s* to accurately model these fast reactions. 2) species R in not modeled as particles.

This is a relatively simple example in which we compute the ODE separately with Python code, however it shows the strength of this approach in which one can couple other physics engines to MCell4 and achieve multi-scale simulations.

## 4 Conclusions

### 4.1 Summary

We have described MCell4, a newly updated particle-based reaction-diffusion tool based on Monte Carlo algorithms that allows spatially realistic simulation of volume and surface molecules in a detailed 3D geometry. MCell4 builds on features of MCell3 (and MCell3-R), providing improved integration with the BioNetGen Language as well as a Python API that enables control of a simulation through Python code.

In MCell4, as opposed to MCell3, molecules and reactions are natively written in BNGL allowing a seamless transition between MCell4 and BNG simulation environments. The update has dramatically improved the ability to run network free simulations in the spatial MCell environment, when compared to the previous MCell3-R which employed the NFSim engine to run reaction written in BNGL [12, 14].

The new Python API, enables one to write Python code that can change geometry, reaction rates, create or remove molecules, execute reactions, etc., during a simulation. This powerful new feature allows construction and execution of multi-scale hybrid models.

As we have demonstrated here through examples, MCell4 adds many new features including the ability to create fully spatial network-free molecular reaction models within realistic geometry. It adds the ability to switch back and forth easily between MCell4 and BNG environments; and it adds the ability to simulate transmembrane or transcellular interactions between surface molecules.

MCell4 is a significant improvement on the previous MCell3-R version with respect to simulation speed, number of features, as well as usability. It allows simulation of new classes of systems that could not be modeled previously.

### 4.2 Availability and Future Directions

MCell4 is available under the MIT license. For easy installation and usage, a package containing MCell, Blender, the Blender plugin CellBlender, and other tools is available along with detailed documentation and on-line tutorials at [37]. MCell4 includes a new C++ library for parsing the BioNetGen language and provides methods to process BioNetGen reactions. This library libBNG is also available under the MIT license [17].

MCell4 does not currently support the definition of spatially extended complexes that could be useful, for instance, when modeling the post-synaptic density [5] or actin filament networks [38] where simply replacing these polymers with a single point in space is inadequate. Furthermore, the ability to model volume exclusion by individual molecules and complexes will be an important goal for the future. We have plans to combine particle-based simulation with concentration or well-mixed simulation algorithms such as SSA [39] or the finite element method that uses PDEs (partial differential equations), e.g., [40]. Such hybrid modeling will provide means to simulate longer timescales while still being spatially accurate and able to correctly handle cases when the copy number of molecules is low. All these features will be the focus of future developments.

## Acknowledgements

The authors thank Dr. Padmini Rangamani for discussions on boundary conditions and the biophysics of diffusion near membranes. We heartily thank Dr. Markus Dittrich, Dr. Burak Kaynak, Dr. Oliver Ernst, Dr. Rex Kerr, Jacob Czech, Jed Wing, Don Spencer, and Robert Kuczewski whose insights on API design for discrete event simulation have guided development of MCell over the years, laying the foundation for MCell4. And we thank Jorge Aldana for his expert technical support of the computing infrastructure in the Computational Neurobiology Laboratory at Salk. Funding for this research was provided by NIH MMBioS P41-GM103712, NIH CRCNS R01-MH115556, NIH CRCNS R01-MH129066, NSF NeuroNex DBI-1707356, and NSF NeuroNex DBI-2014862.

## Notes

https://github.com/mcellteam/article_mcell4_1.git

### Competing Interest Statement

The authors have declared no competing interest.

### Summary of Updates

Revised main text for clarity

